# Branched-chain amino acid assimilation promotes mixotrophy of ammonia-oxidizing archaeal sponge symbionts

**DOI:** 10.1101/2025.09.09.672702

**Authors:** Bettina Glasl, Katharina Kitzinger, Heidi M. Luter, Anton Legin, Erika Salas, Nathalie Heldwein, Katarina Damjanovic, Stefan Schuster, Marie Rutsch, Bram Vekeman, Daan R. Speth, Nicole M. J. Geerlins, Petra Pjevac, Joana Séneca, Margarete Watzka, Wolfgang Wanek, Michael Wagner

## Abstract

Ammonia-oxidizing archaea (AOA) frequently form symbiotic associations with marine sponges. While free-living AOA are generally considered metabolically constrained chemolithoautotrophs, sponge-associated AOA encode for a branched-chain amino acid (BCAA) transporter, suggesting mixotrophic potential. Here, we test the unusual mixotrophic lifestyle of sponge-associated AOA by tracing the assimilation of ^13^C- and ^15^N-labeled BCAA in the sponge holobiont *Ianthella basta*. We demonstrate that BCAA degradation fuels ammonia oxidation and quantify BCAA uptake at the single-cell level by combining stable isotope probing, catalyzed reporter deposition fluorescence *in situ* hybridization, and nanoscale secondary ion mass spectrometry. Our results reveal that sponge-associated AOA are mixotrophic, assimilating BCAA as an additional carbon and nitrogen source. This metabolic adaptation may modulate BCAA availability in the holobiont, potentially regulating the host’s mTOR pathway. Collectively, our study reveals a novel nutritional interaction in sponge holobionts and challenges the perception of constrained metabolic capacities of AOA.

## Introduction

Sponges (Porifera) are one of the oldest extant animal phyla ^1^. These simple filter-feeding invertebrates have evolved in the Precambrian, long before the radiation of all other animal phyla ^2^. Sponges have successfully colonized virtually all aquatic habitats and contribute considerably to the biodiversity of our oceans ^3^. Their unparalleled filtration capacity makes them an important component of many marine ecosystems, where they play a crucial role in the recycling of dissolved organic matter ^4^. Many marine sponges are also known for their highly diverse and abundant microbiomes ^5,6^, whose metabolic functions underpin host health and resilience ^7,8^. The ancestral position of sponges in the evolution of animals and the symbiotic partnership with their metabolically complex microbiomes offer unprecedented insights into the early evolution and fundamental mechanisms of animal-microbe interactions.

Ammonia-oxidizing archaea (AOA) are one of the most widespread and abundant microbiome members of marine sponges ^9,10^. AOA symbionts, often vertically transmitted from the parents to the offspring, play a vital role in the recycling of nitrogenous waste products in the sponge holobiont and have been hypothesized to regulate signaling pathways of their sponge host via nitric oxide production ^11–17^. The symbiotic interaction of AOA with marine sponges is striking because AOA, one of the most successful microbial taxa in the global biosphere ^18^, rarely form symbiotic relationships with animals. Yet, sponge-associated AOA lineages have evolved multiple times within the order *Nitrososphaerales* (formerly known as *Thaumarchaeota) and vary considerably in their host species specificity* ^13,16,19,20^*. Both free-living and sponge-associated AOA* share the *metabolic capacity to derive* energy and reducing equivalents for the fixation of inorganic carbon from the oxidation of ammonia to nitrite. Genomes of sponge-associated AOA seem to support the utilization of alternative nitrogenous waste products for ammonia oxidation, such as urea, nitriles, cyanides, and creatinine ^13,15,16^, a genomic trait shared with free-living AOA ^9^. The utilization of alternative nitrogen sources (i.e., urea, cyanate) for ammonia oxidation has been proven for free-living AOAs ^21,22^; however, experimental evidence for such metabolic versatility in sponge-associated AOA is lacking. In contrast to most free-living AOA, sponge-associated AOA genomes also frequently encode for branched-chain amino acid (BCAA, i.e., valine, leucine, and isoleucine) transporters ^9,13^, which may represent a unique adaptation to their symbiotic lifestyle. Based on the presence of BCAA transporters in sponge-associated AOA genomes, it has been hypothesized that BCAA may serve as alternative carbon and nitrogen sources, thus classifying sponge-associated AOA as mixotrophs rather than strict chemolithoautotrophs ^13^.

Here, we test the hypothesis that sponge-associated AOA can assimilate BCAA as an alternative carbon and nitrogen source. Using the tropical marine sponge *Ianthella basta,* a well-established holobiont model with a highly stable and low-complexity microbiome ^23^, we investigated BCAA uptake by its dominant AOA symbiont, *Candidatus Nitrosospongia ianthellae* ^13^. We reconstructed the genomic potential of both the sponge host and its AOA symbiont to transport, synthesize, and degrade BCAA. To link genomic potential to functional activity, we quantified BCAA-derived ammonia oxidation, inorganic carbon fixation, and BCAA assimilation at bulk and single-symbiont cell levels. The uptake at the single-cell level was quantified using nanoscale secondary ion mass spectrometry (NanoSIMS) in combination with catalyzed reporter deposition fluorescence *in situ* hybridization (CARD-FISH). Our results demonstrate that BCAA are directly used by the AOA symbiont *Ca*. Nitrosospongia ianthellae and likely other sponge-associated AOA lineages in general, confirming the unusual mixotrophic lifestyle of sponge-associated AOA. The ability to take up BCAA may play a critical role in host-symbiont communication by regulating the availability of BCAA, an important signaling molecule in animals.

## Results

### 1) Sponge-associated AOA genera are enriched in BCAA transporters

We conducted a comparative analysis of all publicly available genomes and metagenome-assembled genomes (MAGs; 95% average nucleotide identity, >80% completeness and <5% contamination, Genome Taxonomy Database version 220) of sponge-associated (n = 32), coral-associated (n = 2) and free-living environmental AOA (n = 166) to understand the phylogenetic distribution of BCAA biosynthesis and uptake capabilities across the archaeal family *Nitrosopumilaceae* (Figure 1). All AOA genomes, irrespective of their native habitat or taxonomic affiliation, possess key genes for ammonia oxidation (*amoABC*), carbon fixation via the 3-hydroxypropinate/4-hydroxybutyrate (3-HP/4-HB) cycle (*accABC*), and BCAA biosynthesis via the ilv operon (*ilvABHCDE*). The four distinct sponge-specific AOA genera, *Cenarchaeum* ^20^, *Ca*. Nitrosokoinonia ^16^, *Ca*. Nitrosoabysuss ^19^ and *Ca*. Nitrosospongia ^13^, as well as many other still unnamed sponge-associated AOA, additionally encode the high-affinity LIV-I system (*livKFGHM)*, an ATP-binding cassette (ABC) transporter specific for BCAA ^24^ (Figure 1). Genes encoding for the LIV-I BCAA transport system were predominantly present in sponge-associated AOA (approximately 80% of all sponge-associated AOA genomes analyzed in this study). The majority (93%) of the here analyzed free-living AOA lack known BCAA transport systems; however, we did observe them in 12 genomes from sediment samples, especially from deep-sea and seep sites (representing approximately 7% of all free-living AOA genomes analyzed in this study; see Figure 1), which are often densely colonized by sponges ^25^. The shared evolutionary origin of LIV-I BCAA transporter genes in deep-sea sediment and sponge-associated AOA linages may suggest that the shared habitat has enabled the transfer between those linages.

**Figure 1.**
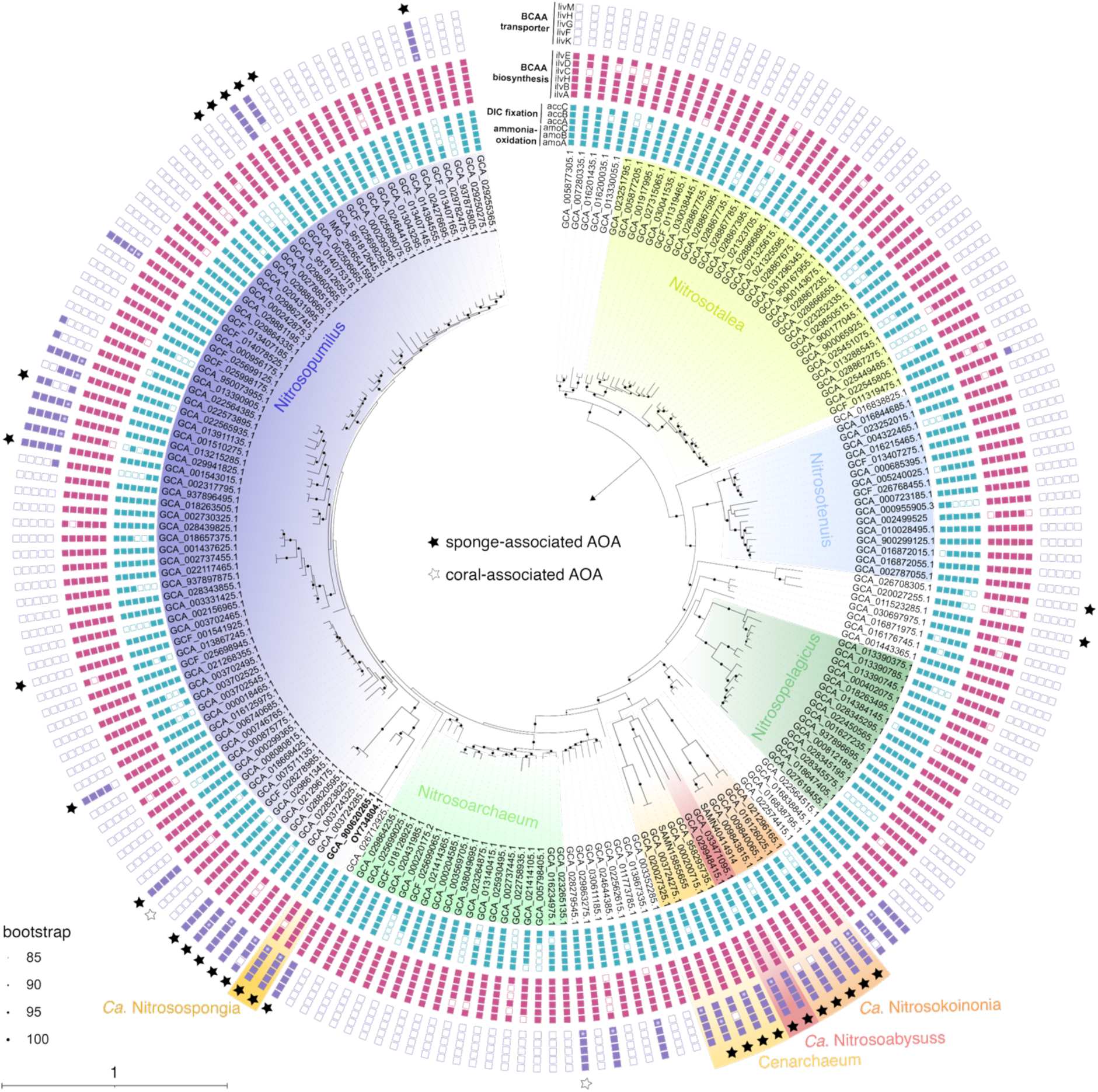
Phylogenomic tree of ammonia-oxidizing archaea. The maximum-likelihood tree of ammonia-oxidizing archaea (AOA) belonging to the family of *Nitrosopumilaceae* (deposited at GTDB version R220) is based on concatenated marker proteins. The tree highlights the presence of marker genes for ammonia oxidation (*amoABC*) and the autotrophic fixation of dissolved inorganic carbon (DIC) via the 3-hydroxypropinate/4-hydroxybutyrate cycle (*accABC*). It further displays the phylogenetic distribution of the branched-chain amino acid (BCAA) biosynthesis gene cluster (*ilvABHCDE*) and the LIV-I BCAA transporter system (*livKFGHM*). Sponge- and coral-associated AOA lineages have evolved multiple times in the *Nitrosopumilaceae family*. Genomes of the sponge-specific AOA genera *Ca*. Nitrosokoinonia, *Ca*. Nitrosoabysuss, *Cenarchaeum*, and *Ca*. Nitrosospongia consistently encode for the LIV-I BCAA transporter. Ultrafast bootstrap (n = 1,000) support is shown on the branches of the tree, and the scale bar corresponds to 1 estimated amino acid substitution per site. A white asterisk (*) inside a colored box indicates genes that were manually annotated. Sponge-associated genomes are indicated with black stars, coral-associated genomes with white stars.

To better understand the evolutionary origins of genes encoding the LIV-I BCAA transport system in *Nitrosopumilaceae,* we performed a phylogenetic assessment of the five subunits (EDF 1). Protein sequences of the LIV-I-BCAA transport system appear to be monophyletic, with the exception of *livF* (K01996), despite the fact that sponge-associated AOA lineages have evolved multiple times within this family. The phylogenetic neighborhood of the *livGFHM* genes (K01995-k01998) consists largely of other members of the *Thermoproteota*, including the members of the early-branching lineages of the putative chemoorganoheterotrophic *Nitrososphaerales* order that lack the ability to oxidize ammonia and fix inorganic carbon ^26^. However, the phylogeny of these genes is inconsistent and additionally incongruent with (concatenated) marker gene phylogenies of the *Thermoproteota* phylum. In addition, the *livK* gene of the LIV-I-BCAA transport system is closely related to *Bacteroidota* sequences, and thus appears to have a different evolutionary history from the rest of the complex, despite occurring in an operon in the AOA. It thus seems likely that the *Nitrosopumilaceae* received four of their *liv*-genes through horizontal gene transfer from another member of the *Thermoproteota* phylum, and acquired *livK* separately. Subsequently, it is likely that several transfers within the *Nitrosopumilaceae* spread the *liv*-genes to sponge-associated and (potentially) free-living AOA. Overall, the striking association of BCAA uptake systems with sponge-associated AOA indicates that this capability confers an important adaptation to their symbiotic lifestyle within sponges.

### 2) Genomic potential of the *I. basta* sponge and its AOA symbiont to utilize BCAA

To better understand the role of BCAA in the AOA-sponge symbiosis, we analyzed in detail the respective genomes of the AOA symbiont *Ca*. Nitrosospongia ianthellae (GCA_963457515.1) and its sponge host *Ianthella basta* (GCA_964019415.1). The AOA symbiont *Ca*. N. ianthellae comprises, based on 16S rRNA gene amplicon data, on average 55.2% (SD 10.6%) of the *I. basta* microbiome (EDF 2), and its genome encodes for all hallmark genes for chemolithoautotrophic ammonia oxidation and inorganic carbon fixation (Figure 2). In addition, the *Ca*. N. ianthellae genome encodes for the biosynthesis of BCAA (*ilv* and *leu* gene cluster) and the LIV-I BCAA transport system (Figure 2). The latter has previously been found to be expressed in *Ca*. N. ianthellae in the holobiont using metaproteomics ^13^. Furthermore, *Ca*. N. ianthellae has the genomic potential to perform the first step of the BCAA degradation pathway, the transamination of BCAA using a BCAA aminotransferase (*ilvE*), resulting in the production of glutamate and branched-chain keto acids (BCKA). Deamination of glutamate using the encoded glutamate dehydrogenase leads to the production of ammonium and thus provides an alternative way for providing the substrate for the energy metabolism of this AOA. The second step of the BCAA degradation pathway is the decarboxylation of the BCKA, catalyzed in other organisms by the BCKA dehydrogenase complex (EC 1.2.4.4) or by the BCKA ferredoxin reductase (EC 1.2.7.7). Homologous genes encoding for both enzymes are missing from the *Ca*. N. ianthellae genome. Yet, the *Ca*. N. ianthellae genome encodes for a member of the multi-enzyme family 2-oxoacid:ferredoxin oxidoreductase (Figure 2). Enzymes of this family mediate the decarboxylation of pyruvate, 2-oxoglutarate, and BCKA to their corresponding acyl-CoA derivatives ^27–29^, indicating 2-oxoacid:ferredoxin oxidoreductase may be involved in BCAA degradation in *Ca.* N. ianthellae and other AOA.

**Figure 2.**
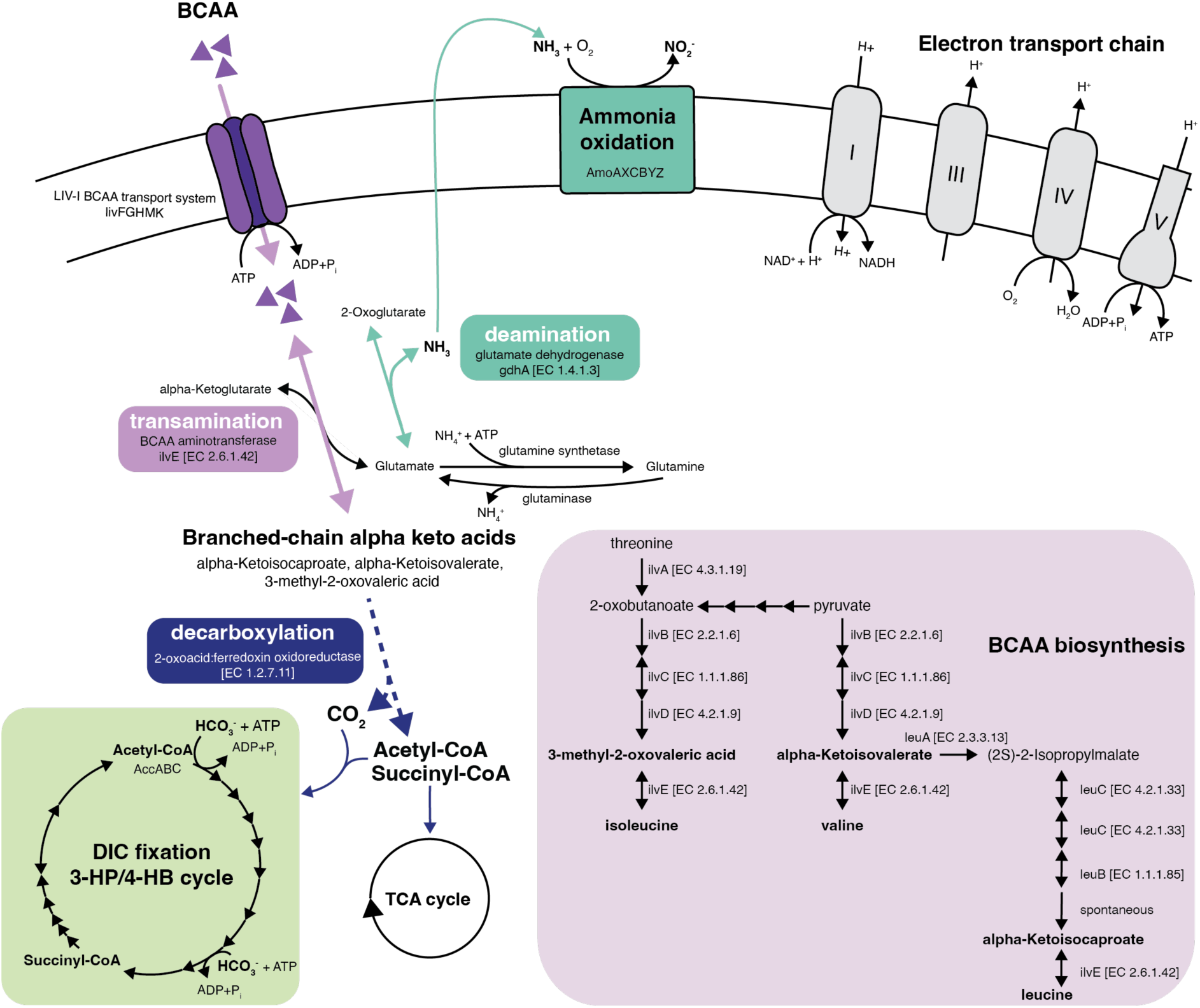
Branched-chain amino acid (BCAA) degradation and biosynthesis pathways of the ammonia-oxidizing archaeal symbiont *Candidatus* Nitrosospongia ianthellae. The *Ca*. N. ianthellae genome encodes for the high-affinity LIV-I branched-chain amino acid (BCAA, i.e., leucine, valine, and isoleucine) transport system. The first step in BCAA degradation is the transamination of BCAA with alpha-ketoglutarate. The enzyme branched-chain aminotransferase catalyzes this reversible reaction, transferring an amino group from the BCAA to alpha-ketoglutarate and creating glutamate and the corresponding branched-chain alpha keto acid (BCKA). Glutamate can then be used for protein synthesis, amino acid biosynthesis, and glutamine formation. Alternatively, glutamate can be deaminated by the glutamate dehydrogenase enzyme, releasing ammonia (NH_3_), which can be used for ammonia oxidation. In the second step of the BCAA degradation, BCKA are decarboxylated either by an unknown branched-chain alpha keto acid dehydrogenase complex (BCKDC) (not shown) or by 2-oxoacid:ferredoxin oxidoreductase, thereby releasing CO_2_ and producing acetyl-CoA, and succinyl-CoA. These molecules can enter both the dissolved inorganic carbon (DIC) fixation pathway and the tricarboxylic acid (TCA) cycle. In addition, *Ca*. N. ianthelleae encodes all the hallmark genes characteristic of the chemolithoautrophic lifestyle of ammonia-oxidizing archaea, including the ammonia-monooxygenase enzyme (AmoAXCBYZ), the electron transport chain (complex I-V), and the 3-hyroxypropionate/4-hydroxybutyrate (3-HP/4-HB) cycle for DIC fixation. The dashed line represents a hypothetical pathway.

In contrast to its AOA symbiont, the sponge *I. basta* lacks the genomic repertoire for BCAA biosynthesis but possesses the complete set of genes encoding for the BCAA degradation pathway (EDF 3a). Furthermore, the *I. basta* genome also encodes for the BCAA-stimulated mTOR complex 1 (mTORC1, EDF 3b), which is part of the conserved mechanistic (formerly mammalian) target of rapamycin (mTOR) pathway in animals ^30^. Homologous leucine sensors of the mTORC1 pathway were not detected in the *I. basta* genome (EDF3b), and further investigations are needed to better understand the regulatory role of BCAA on the mTORC1 pathway in sponges. The mTORC1 pathway in sponges remains understudied, but it has been shown that mTORC1 plays an important role in integrating symbiont-derived nutrients into host metabolism and homeostasis in cnidarians ^31^, deep-sea mussels ^32^, and aphids ^33^. The effect of BCAA on the microtubule organization via mTORC1 (EDF 3b) may also have a vital role in the functioning and formation of ciliated cell types in sponges such as the choanocytes, a highly proliferative cell type that contributes to tissue regeneration, sexual reproduction, and is specialized to capture food particles ^34,35^. Overall, our analysis suggests that *I. basta* is auxotrophic for BCAA and requires the active uptake of BCAA. As such, the ability of its AOA symbiont to modulate the BCAA availability in the sponge holobiont may play a fundamental role in the host-symbiont communication.

### 3) BCAA use by the *Ianthella basta* sponge holobiont

The *I. basta* pore water contained substantial concentrations of free BCAA — 2.19 ± 0.38, 1.97 ± 0.75, and 0.81 ± 0.13 µmol L⁻¹ for valine (Val), leucine (Leu), and isoleucine (Ile), respectively, measured in field control samples (EDF 4a). This indicates that both the sponge and its symbionts are consistently exposed to BCAA, which may serve as potential substrates. To test whether BCAA are actively utilized by the holobiont, we conducted 24h incubation experiments using an equimolar mix of fully ^13^C- and ^15^N-labeled Val, Leu, and Ile (1 µmol L⁻¹ total BCAA concentration), with parallel incubations containing either sponge explants or filtered seawater (EDF 5).

Over the course of the incubation, the labeled BCAA were progressively depleted, with slightly faster consumption observed in incubations containing sponge explants (EDF 4b). Correspondingly, the sponge holobiont biomass showed slight enrichment in both ^13^C and ^15^N after 24 h compared to non-incubated sponge field controls (EDF 4c), indicating active assimilation of BCAA-derived carbon and nitrogen by the holobiont. Alongside BCAA depletion, we detected increasing levels of dissolved ^13^C-labeled CO₂ in the incubation seawater, reaching up to 4 µmol L⁻¹ (EDF 4d). Given that complete respiration of the added BCAA would produce approximately 5.6 µmol L^-1^ ^13^C-CO₂, this suggests that a substantial fraction (approximately 70%) of the supplied BCAA was respired.

As a first indication of whether sponge-associated AOA may contribute to the observed BCAA use, we assessed whether some of the BCAA-derived nitrogen was nitrified to ^15^N-nitrite. *I. basta* contains no nitrite-oxidizing symbiont ^13^, therefore nitrite does not get further oxidized to nitrate. Indeed, we observed significant and linear production of both isotopically unlabeled and ^15^N-labeled nitrite in the incubations containing sponge explants (Figure 3, EDF 6). This proves that the symbiotic AOA of *I. basta* were both metabolically active and were using BCAA-derived ^15^N-labeled nitrogen, with up to 56% of the provided ^15^N from BCAA being recovered as ^15^N-nitrite.

**Figure 3.**
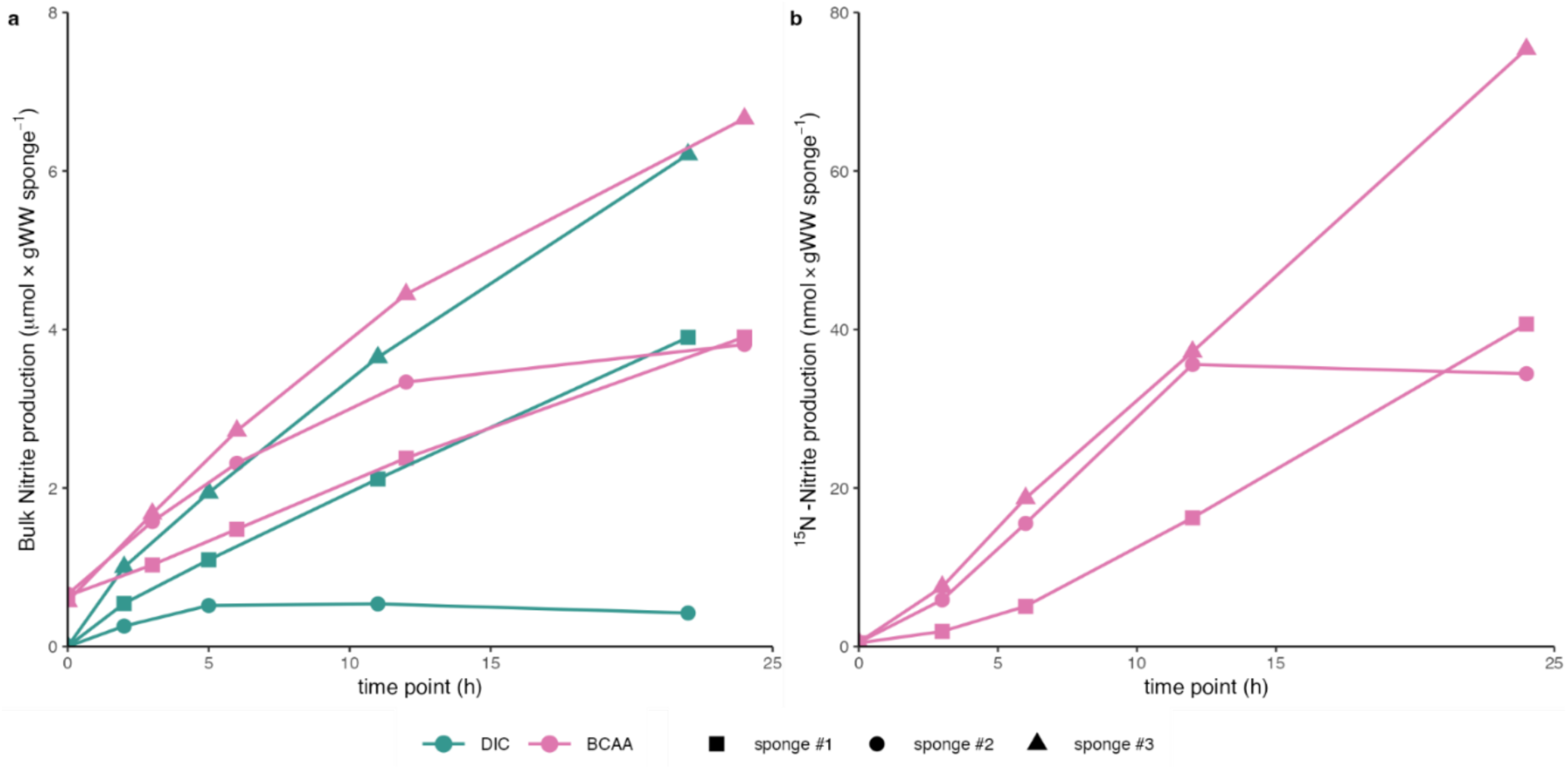
Nitrification activity of the *Ianthella basta* sponge holobiont. a) Bulk nitrite (NO_2_^-^) production per sponge individual during ^13^C-labeled dissolved inorganic carbon (DIC) and ^13^C^15^N-labeled branched-chain amino acid (BCAA) incubation experiments. Nitrite formation in each incubation jar was normalized to the sponge wet weight (ww) of the sponge explants. No nitrite formation was detected in the seawater control jars (see EDF 6a). b) ^15^N-nitrite (^15^N-NO_2_^-^) production per sponge individual in the BCAA incubation experiment. Sponge explants were incubated in filtered seawater amended with an equimolar mix of ^13^C^15^N-labeled valine, leucine, and isoleucine. The ^15^N-nitrite formation in each incubation jar was normalized to the sponge wet weight (ww) of the sponge explants. No nitrite formation was detected in the seawater control jars (see EDF 6b).

However, these bulk analyses cannot differentiate between direct and indirect use of BCAA for nitrification by the symbiotic AOA. The heterotrophic bacterial symbionts, *Ca*. Luteria ianthellae and *Ca*. Taurinisymbion ianthellae, of the *I. basta* holobiont have the genetic capacity for BCAA degradation ^23^ and thus can release ^15^N-labeled ammonium, which can subsequently be oxidized to nitrite by the symbiotic AOA.

### 4) Symbiotic AOA cells assimilate both BCAA and ^13^C-CO_2_

To determine whether *Ca.* N. ianthellae is capable of directly taking up BCAA and utilizing these amino acids for growth; we measured ^13^C- and ^15^N-assimilation from BCAA into single symbiont cells. To achieve this, we specifically visualized the three dominant symbionts of *I. basta*, the AOA symbiont (*Ca.* N. ianthellae), the alphaproteobacterial symbiont *Ca.* Luteria ianthellae, and the gammaproteobacterial symbiont *Ca.* Taurinisymbion ianthellae, using catalyzed-reporter deposition fluorescence *in situ* hybridization (CARD-FISH) in combination with DAPI (4’,6-diamidino-2-phenylindol) staining and measured cellular isotopic enrichment by nanoscale secondary ion mass spectrometry (NanoSIMS). While isotopic enrichment was negligible in the measured sponge biomass, all symbiont groups were significantly enriched in both ^13^C and ^15^N after 24 h incubation with ^13^C^15^N-labeled BCAA. This experimental evidence is consistent with the genomic capacity of all three main symbionts to import and utilize BCAA.

The highest isotopic enrichment for both ^13^C and ^15^N was observed in the gammaproteobacterial symbiont cells (1.60 ± 0.37 and 0.88 ± 0.43, mean ± sd ^13^C-at% and ^15^N-at% enrichment, respectively), followed by alphaproteobacterial symbiont cells (1.37 ± 0.16 and 0.65 ± 0.22) and the AOA symbiont cells (1.19 ± 0.08 and 0.42 ± 0.09; Figure 4). This pattern may reflect generally higher cellular activity in the heterotrophic gamma- and alphaproteobacterial symbionts compared to symbiotic AOA *Ca.* N. ianthellae. The mean ratio of new carbon to nitrogen assimilation, considering significantly enriched cells, was similar for the gammaproteobacterial and alphaproteobacterial symbionts (10.5 ± 8.8 and 10.1 ± 6.8 fmol-C cell⁻¹ day⁻¹ to fmol-N cell⁻¹ day⁻¹, respectively), whereas the symbiotic AOA *Ca.* N. ianthellae assimilated relatively more carbon (16.9 ± 5.5 fmol-C cell⁻¹ day⁻¹ to fmol-N cell⁻¹ day⁻¹).

**Figure 4.**
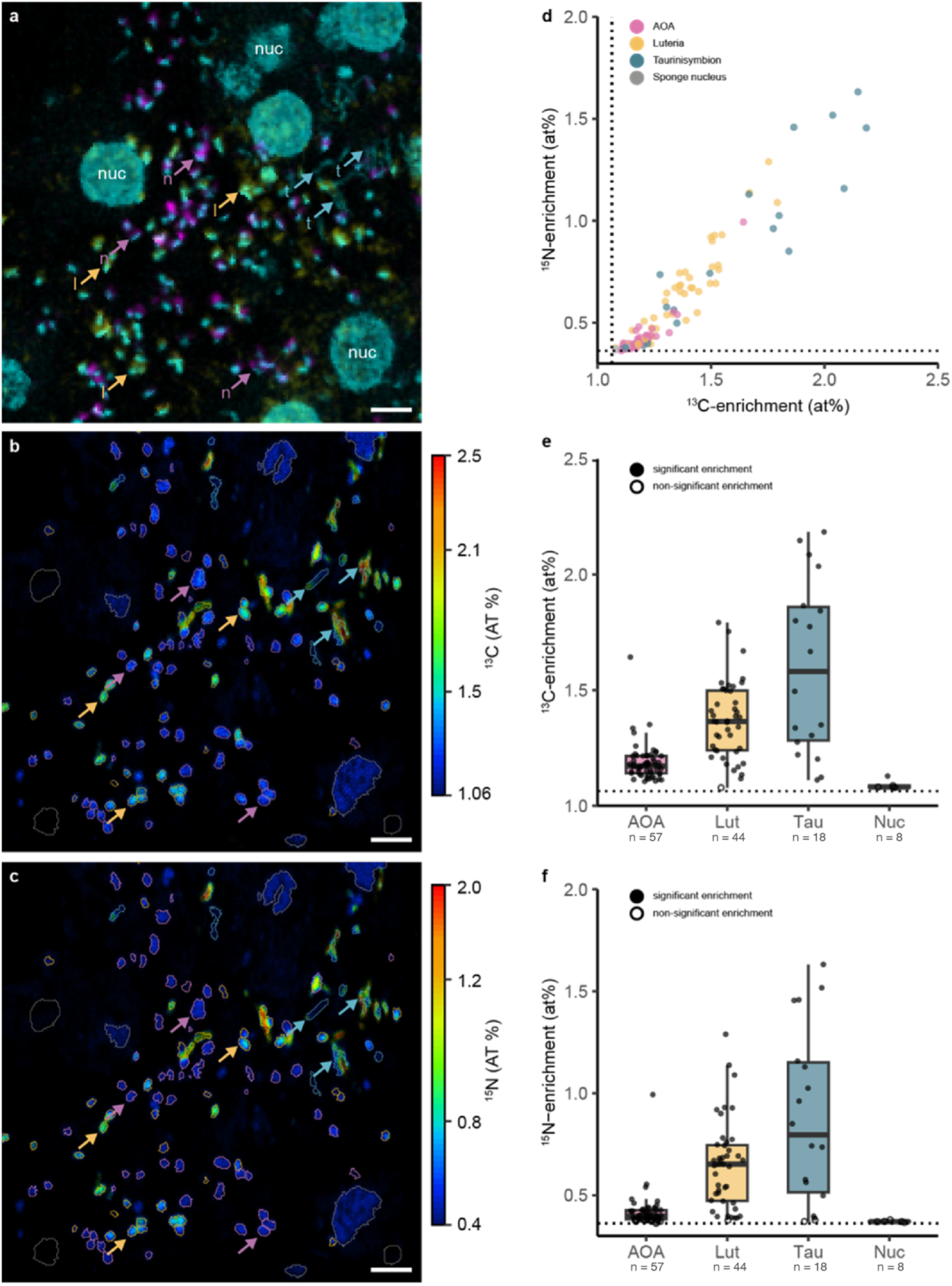
Single-cell ^13^C and ^15^N-assimilation from branched-chain amino acids (BCAA) measured by NanoSIMS. a) CARD-FISH image counterstained with DAPI, visualizing the ammonia-oxidizing archaeal symbiont *Ca*. Nitrosospongia ianthellae (n, magenta, stained by probe Arch915), the alphaproteobacterial symbiont *Ca*. Luteria ianthellae (l, yellow, stained by probe AlfD729), the gammaproteobacterial symbiont *Ca*. Taurinisymbiont ianthellae (t, cyan), and the sponge nucleus (nuc, cyan, counterstained with DAPI) in a 5 µm cryosection of the *Ianthella basta* holobiont. b-c) Corresponding NanoSIMS images of ^13^C at% enrichment and ^15^N at% enrichment after the incubation with isotopically labeled BCAA. Quantified regions of interest of the dìerent cell types are marked by colored outlines (*Ca*. N. ianthellae in magenta, *Ca*. L. ianthellae in yellow, *Ca*. T. ianthella in cyan, sponge nucleus and tissue marked in grey). The scale bar is 2 µm in all images. The saturation intensity (hue) of enrichment was modulated by the measured phosphorus signal for better visibility. d) Correlation between ^13^C and ^15^N single cell isotopic enrichment from ^13^C^15^N-labeled BCAA incorporation. e-f) Single cell ^13^C- and ^15^N-enrichment of the main *I. basta* microbial symbiont groups (AOA = *Ca.* N. ianthellae, Lut = alphaproteobacterial *Ca*. Luteria ianthellae, Tau = gammaproteobacterial *Ca.* Taurinisymbion ianthellae) and sponge nuclei (Nuc). Boxplots depict the 25– 75% quantile range, with the center line depicting the median (50% quantile); whiskers encompass data points within 1.5× the interquartile range. Dotted lines in d-f indicate ^13^C and ^15^N natural abundance at% as measured for symbiont cells in non-incubated sponge biomass (1.06 ^13^C at% and 0.36 ^15^N at%, respectively). The total number of measured cells within each group is indicated below the x-axis labels.

The consistent enrichment in both ^13^C and ^15^N of the symbiotic AOA *Ca.* N. ianthellae provides strong evidence for their direct BCAA utilization. Any ^13^C-DIC derived from heterotrophic breakdown of BCAA would have been immediately diluted in the large ambient DIC pool of natural isotopic composition (seawater contains approx. 2.3 mM DIC; we anticipate similar concentrations in the sponge mesohyl). Thus, only minimal ^13^C-labeling of *Ca.* N. ianthellae from inorganic carbon fixation due to cross-feeding would be expected. Likewise, ^15^N-ammonium released from heterotrophic breakdown of ^13^C^15^N-labeled BCAA would also be diluted in the ambient ammonium pool (from, e.g., organic matter breakdown by the sponge and/or its symbionts; EDF 7), lowering the likelihood of ^15^N cross-feeding from BCAA.

To date, autotrophic inorganic carbon fixation by *Ca.* N. ianthellae has been inferred from the correlation between nitrification activity and bulk ^13^C-DIC uptake by the sponge holobiont ^13^, as well as from the presence of the 3-HP/4-HB carbon fixation pathway in their genomes and the metaproteome ^13,23^. However, single-cell evidence for autotrophic C-fixation by *Ca.* N. ianthellae within the sponge holobiont has been lacking. To test whether *Ca.* N. ianthellae also actively fix inorganic carbon, in addition to BCAA utilization, we conducted parallel incubations with ^13^C-DIC and measured isotope incorporation at the single-cell level using NanoSIMS. In these incubations, as in the BCAA incubations, sponge-associated AOA were metabolically active, as evidenced from increasing nitrite concentrations (Figure 3, EDF 6), and *Ca.* N. ianthellae fixed significantly more ^13^C-DIC than other microbial cells in the holobiont (one-sided Mann– Whitney U test, W = 14,007, p < 2.2 × 10⁻¹⁶; EDF 8a).

Using the measured carbon enrichment from both BCAA and DIC for *Ca.* N. ianthellae, we estimated single-cell C-based growth rates in our incubations to be 0.011 ± 0.007 and 0.018 ± 0.010, respectively. These correspond to doubling times of ∼95 and ∼55 days, respectively, from BCAA and DIC. Such long doubling times are unusual compared to other marine AOA, e.g., 0.7 days in pure culture ^36^ to several days for free-living marine AOA in the eutrophic Gulf of Mexico ^37^. The slow growth rates observed here may indicate that sponge-associated AOA require additional C-sources in addition to CO2 and BCAA. However, further experiments are needed as the slow growth rates may also reflect the incubation conditions, dilution of cellular carbon by the CARD-FISH procedure (see Materials and Methods), or alternatively, an inherently slow turnover of the AOA population within the sponge holobiont. Nevertheless, our measurements demonstrate that BCAA assimilation can contribute substantially (ca. 58% of autotrophic C-fixation) to *Ca.* N. ianthellae growth in the sponge holobiont under our incubation conditions (EDF 8b). Notably, this estimate assumes that symbiont cells are exposed both to the isotope concentrations supplied to the incubation seawater and to the measured BCAA concentrations in sponge pore water. A more conservative assumption, assuming that symbionts do not experience pore-water BCAA concentrations but only the labeled BCAA provided, still yields BCAA-derived growth equivalent to ∼10% of autotrophic C-fixation-derived growth (EDF 8b).

Taken together, our results show that the AOA symbiont *Ca.* N. ianthellae assimilates both carbon and nitrogen from BCAA in the sponge holobiont, in addition to autotrophic C-fixation, thereby supporting a mixotrophic lifestyle. Beyond potentially regulating host physiology through BCAA level modulation, the symbiotic AOA *Ca.* N. ianthellae benefit from organic compound assimilation to enhance growth rates.

## Discussion

Ammonia-oxidizing archaea are generally considered to be chemolithoautotrophs, yet there has been a long-standing debate over whether they are obligate autotrophs or can grow mixotrophically ^38–42^. The observation that alpha-keto organic acids, such as pyruvate, stimulated the growth of some AOA pure cultures ^43,44^ initially suggested a mixotrophic lifestyle of some of these ammonia-oxidizer strains. However, later it was found out that alpha-keto organic acids are not assimilated as a carbon source during ammonia oxidation but instead stimulate AOA growth via the non-enzymatic detoxification of H_2_O_2_ ^45^, which can inhibit catalase-negative AOA. Amino acid use by AOA has also been tested extensively. While pure culture cultures of the marine AOA *Nitrosopumilus adriaticus* and *Nitrosopumilus piranensis*, which do not encode a BCAA uptake system, did not use leucine as a source of energy and/or carbon ^46^, early studies showed incorporation of label from a mixture of 15 tritiated amino acids ^47^ and tritiated leucine ^48^, respectively, into marine free-living AOA (not yet recognized as such in those studies) cells using microautoradiography-FISH. Despite the fact that some marine free-living AOA are encoding BCAA uptake systems (Figure 1), the label incorporation observed in these former studies could also stem from cross-feeding from other community members that were the primary users of the labeled amino acids. More recent single-cell analyses of environmental marine AOA communities using a ^13^C^15^N-labeled algal amino acid mixture pointed toward the use of amino acid-derived nitrogen, likely via cross-feeding ^49^. Taken together, while amino acid labeling experiments with marine free-living AOA demonstrated some activity, no direct evidence for mixotrophic growth of these nitrifiers could yet be provided.

Here, we show for the first time the simultaneous assimilation of ^13^C and ^15^N from BCAA by an AOA by analyzing the sponge-associated AOA *Ca*. N. ianthellae *in situ* using CARD-FISH - NanoSIMS (Figure 4). We also experimentally demonstrate the concurrent capacity of *Ca*. N. ianthellae for fixation of dissolved inorganic carbon (EDF 8a) and thus provide compelling evidence for a mixotrophic lifestyle of these ammonia-oxidizers in the sponge holobiont. As the *de novo* biosynthesis of BCAA is energetically costly for cells and ranges from 10 ATP per molecule of valine, 4 ATP per molecule of leucine, and 3 ATP per molecule of isoleucine ^50^, the ability to take up BCAA from the environment, in addition to their *de novo* biosynthesis, might confer a selective advantage for sponge-associated AOA lineages. This is particularly relevant given that BCAA concentrations in the sponge pore water samples (EDF 4) exceeded those previously reported for seawater by 2-3 orders of magnitude ^51^. Simultaneous BCAA uptake via the LIV-I transport system and autotrophic carbon fixation via the 3-HP/4-HB cycle appears to be a widespread adaptation among sponge-associated ammonia-oxidizing archaea (Figure 1). The uptake of BCAA by sponge-associated AOA seems to be a fundamental adaptation to their symbiotic lifestyle. Despite the independent evolution of distinct sponge-associated AOA lineages, subunits of the BCAA LIV-I transporter system are of monophyletic origin (EDF 1). Whether the uptake of BCAA LIV-I transporter subunits into sponge-associated AOA genomes stems from joint inheritance or is a result of joint loss by most free-living AOA cannot be conclusively determined from our analyses. The monophyly of the AOA BCAA transporter genes points to a single acquisition within the AOA. In addition, all BCAA LIV-I transporter-containing AOA share a common ancestor within the *Nitrosopumilaceae*, and therefore, the distribution of the transporter can be explained by one acquisition and multiple losses. However, the BCAA gene phylogenies are incongruent with the genome phylogeny. Notably, the deepest branching BCAA-containing clade in the genome phylogeny (DRGT01) is among the most derived groups in the gene phylogenies. This pattern is more consistent with a single ancestral acquisition followed by frequent horizontal gene transfer among AOA lineages. While the few free-living AOA genomes that encode a LIV-I BCAA transport system (e.g., deep-sea sediment clade DRGT01) share an evolutionary origin with the sponge-associated AOA, the uptake of LIV-I BCAA genes seems to have originally occurred in a sponge-associated AOA.

The sponge *I. basta*, like all animals, is auxotrophic for BCAA and relies on the uptake of BCAA (EDF 3). The readily available BCAA pool in *I. basta* can derive from internal sources, such as phagocytosis of symbionts, the secretion of BCAA by symbionts into the mesohyl, and from the surrounding environment via intercellular gaps between sponge cells ^52^ or the engulfment of dissolved organic matter via pinocytosis ^53,54^. Similar to its ammonia-oxidizing archaeal symbiont, we were able to show that also the heterotrophic symbionts of *I. basta* take up exogenously supplied BCAA as a carbon and nitrogen source (Figure 4) despite being genomically equipped to biosynthesize their own BCAA ^23^. Thus, all three symbionts of *I. basta* have the ability to modulate BCAA concentrations, potentially regulating the availability of essential amino acids in their auxotrophic host (Figure 5). BCAAs may play a role in the biosynthesis of bioactive secondary metabolites, such as bastadins, a class of bromotyrosine-derived macrolactams, which are highly abundant in the *I. basta* holobiont ^55,56^. Bastadins are likely assembled via non-ribosomal peptide synthetase pathways that incorporate amino acid precursors, potentially including BCAAs or their alpha-keto acids ^55^. This suggests that the availability of BCAAs in the sponge holobiont could directly influence the production of these compounds, linking amino acid metabolism to the sponge’s chemical defence arsenal and the generation of molecules with pharmacological potential. Furthermore, BCAA, such as leucine, are also known regulators of the mTORC1 pathway, a conserved pathway in eukaryotes ^30^. Besides its central role in cell proliferation, autophagy, and protein synthesis, mTORC1 also has a functional role in various nutritional symbioses, including the coral-algal ^31^, the aphid-buchnera ^33^, and the mussel-bacteria symbiosis ^32^. In later, mTORC1 regulates phagosome digestion of symbiosome in *Bathymodiolus* mussels in response to the nutrient supply from the symbionts ^32^. However, whether mTORC1 plays a similar role in sponges and whether microbial symbionts actively influence this central eukaryotic pathway by altering BCAA levels remains unknown, in particular, as in the *I. basta* genome the sensing proteins Sestrin2 and Sar1B could not be identified (also lacking in the genome of the sponge *A. queenslandica*, data not shown). Future studies targeting the interaction between symbiont metabolism and host mTOR pathways will be key to gaining insights into the regulatory mechanisms in sponge holobionts.

**Figure 5.**
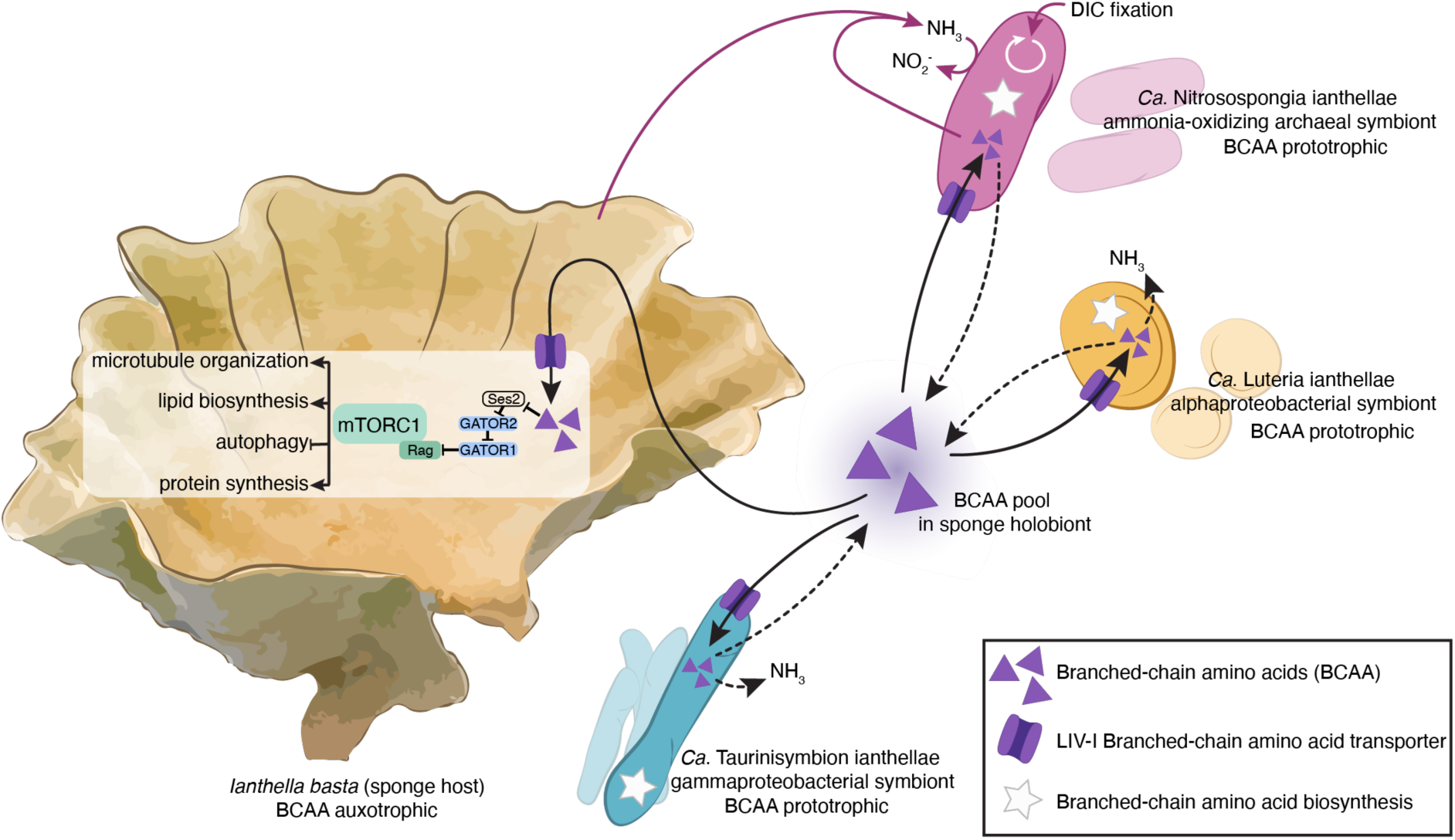
Schematic overview of branched-chain amino acid (BCAA) biosynthesis and assimilation capabilities in the *Ianthella basta* holobiont. The three dominant symbionts of the sponge *I. basta*, the archaeal *Ca*. Nitrosospongia ianthellae, the alphaproteobacterial *Ca*. Luteria ianthellae and the gammaproteobacterial *Ca*. Taurinisymbion ianthellae, take up BCAA (valine, leucine, and isoleucine) as carbon and nitrogen source from the environment (exogenous supply of BCAA). Additionally, all three symbiont genomes possess a full set of genes that encode BCAA biosynthesis. Unlike the two heterotrophic bacterial symbionts, the ammonia-oxidizing archaeal symbiont can also fix dissolved inorganic carbon (DIC) autotrophically with energy derived from the oxidation of ammonia (NH_3_), a metabolic waste product of the sponge host and a product of BCAA catabolism. Thus, the ammonia-oxidizing archaeal symbiont derives its carbon for growth from autotrophic and heterotrophic modes of nutrition (mixotrophy). The BCAA pool in the sponge holobiont is potentially fueled by exogenous BCAA uptake, the phagocytosis of microbial cells, and the BCAA biosynthesis of symbiont cells. The sponge host itself lacks the genomic repertoire for BCAA biosynthesis and therefore relies on the uptake of these essential amino acids from the BCAA pool. BCAA are known to stimulate in other animals the highly conserved mTORC1 signaling pathway, which is fully encoded in the *I. basta* genome (although known leucine-sensing systems are not annotated); thus, BCAA regulation by sponge-symbionts may play a crucial role in host-microbe interactions in this basal metazoan phylum.

## Materials and Methods

### Phylogenetic distribution of BCAA uptake and biosynthesis genes in the ammonia-oxidizing archaeal family *Nitrosopumilaceae*

Metagenome assembled genomes (MAGs; 95% average nucleotide identity, >80% completeness and <5% contamination) of the family *Nitrosopumilaceae* included in the Genome Taxonomy Database (GTDB version R220) and MAGs of publicly available sponge-associated AOA (99% average nucleotide identity, >80% completeness and <5% contamination) were used for the comparative genome analysis. In total, 200 MAGs were included in the analysis, of which 32 MAGs originated from marine sponges and 2 MAGs from corals. Concatenated amino acid sequences of marker proteins were obtained using checkM v1.2.3 ^57^ and used for phylogenomic reconstruction with IQ-TREE v2.4.0 ^58^ using the best-fit model (Q.insect+I+R8) according to the Bayesian Information Criterion (BIC) and 1,000 ultrafast bootstraps. Genomes of *Nitrososphaera viennensis* and *N. gargensis* were used as the outgroup. The phylogenetic tree was subsequently visualized in iTOL v7 ^59^, highlighting sponge- and coral-associated AOA symbiont linages, the presence of genes encoding for LIV-I BCAA transporter, BCAA biosynthesis, and key genes for ammonia-oxidation (*amoABC*) and carbon-fixation (*accABC*) annotated using eggnog mapper v2.1.10 ^60^.

Proteins annotated as K01995 (*livG*), K01996 (*livF*), K01997 (*livH*), K01998 (*livM*), and K01999 (*livK*), corresponding to the 5 subunits of the LIV-I BCAA transport system, were retrieved from the GlobDB release 220 ^61^. Length distribution of the annotated proteins was assessed using “seqkit watch” ^62^, and likely false positives were removed by restricting the hits to proteins of length ranges 230-280aa (K01995), 220-280aa (K01996), 275-320aa (K01997), 260-460aa (K01998), and 350-470aa (K01999). False negatives in the KEGG annotation, and proteins in newly available genomes, were subsequently added by using each curated protein set as seed in a protein alignment score ratio analysis against the GlobDB release 226 ^63^, resulting in final datasets of 371,412 (K01995), 439,115 (K01996), 413,331 (K01997), 393,125 (K01998), and 585,274 (K01999) amino acid sequences. Sequence datasets were clustered at 50% sequence identity (with the exception of K01999, which was clustered at 40% identity) using usearch v12.0 ^64^, aligned using muscle5 v5.3 ^65^, and trees were calculated using FastTree2 v2.1.11 ^66^.

The presence of AOA BCAA genes in the phylogenies described above was assessed in iToL v7 ^59^, and for all subunits the AOA genes clustered in a single clade of the full phylogeny. Several singletons clustered outside this clade of interest but were determined to likely be binning errors after inspection. For the clade of interest of each gene, trees containing all sequences, without prior identity clustering, were calculated as described above. For three subunits (K01995, K01997, K01998), the AOA genes formed a monophyletic clade, whereas for the remaining two subunits (K01996, K01999) two distinct clades were observed.

#### Metabolic reconstruction of Ca. Nitrosospongia ianthellae and Ianthella basta

The closed genome of the ammonia-oxidizing archaeal symbiont *Ca*. Nitrosospongia ianthellae (GCA_963457515.1) was annotated using eggnog mapper v2.1.10 ^60^. Annotations were used to manually reconstruct the following key metabolic pathways: ammonia oxidation, electron transport, inorganic carbon fixation via the 3-hyroxypropionate/4-hydroxybutyrate (3HP/4HB) cycle, biosynthesis of BCAA, BCAA transporter, and BCAA degradation. The *livK* gene encoding for the substrate binding subunit of the LIV-I BCAA transporter failed to annotate using eggnog-mapper. We therefore screened the protein sequences of the *Ca*. N. ianthellae genome for LIV-K protein family (PF13458) using the hmmscan function of HMMER v3.4 (hmmer.org). The *livK* gene is located next to the other LIV-I transporter genes in the *Ca*. N. ianthellae genome.

The *Ianthella basta* genome (GCA_964019415.1; 215M bp, haploid number = 5, chromosome number = 16, genome completeness = 71.6%), previously sequenced and annotated with BRAKER2 ^67^ by the Aquatic Symbiosis Genomics Project ^68^, was submitted to the GhostKOALA platform for automatic KO assignment and KEGG pathway reconstruction ^69^. The completeness of the BCAA biosynthesis pathway, the BCAA degradation pathway, and the mTOR pathway was assessed, and pathway maps were reconstructed using KEGG Mapper.

### Sponge collection

Three individuals of the sponge species *Ianthella basta* (yellow, thin morphotype) were collected from Davies Reef (18°49’19.1"S 147°38’11.3"E, Great Barrier Reef, Australia) at 11 m depth in December 2022. Sponges were transferred to the flow-through aquaria system at the National SeaSimulator at the Australian Institute of Marine Science (Townsville, Australia). After a two-week acclimation period, each sponge individual (n = 3) was fragmented into three equal-sized explants of the fan-shaped sponge body (approximately 5 cm x 7 cm) using sterile scissors. One of the three explants per sponge individual (n = 3) was subsampled for amplicon sequencing and BCAA pore water concentrations. The other two explants of each sponge individual (total n of explants = 6) were kept in an outdoor flow-through aquaria system (2,500 L) under natural lighting and ambient seawater temperatures (approximately 28.5°C) for one week prior to further experiments. An additional *I. basta* individual was collected at the same sampling site in March 2024 as a natural abundance control and for amplicon sequencing. All samples were collected under the permit G21/38062.1 issued by the Great Barrier Reef Marine Park Authority.

### Stable isotope incubation experiments of *Ianthella basta* explants

Custom-made sponge incubation jars (n = 9; EDF 5) were acid-washed using 1M HCl, rinsed, and filled with 800 mL of 0.2 µm filter-sterilized seawater (Sterivex, Merck Pty Ltd), and sealed to avoid exchange with the atmosphere. Six of the jars were amended with an isotopically labeled BCAA mix (1 µM total concentration, L-valine-^13^C_5_^15^N, L-leucine-^13^C_6_^15^N, and L-isoleucine-^13^C_6_^15^N each at 0.33 µM concentration) and 5 µM ^14^NO_2_^-^ as a trap for any produced ^15^N-NO_2_^-^ from BCAA-derived ammonia oxidation. The remaining three jars were amended with 500 µM NaH^13^CO_3_ (dissolved inorganic carbon, DIC). One explant from each sponge individual (n = 3) was placed in one of the two different stable isotope experiments: i) BCAA incubation and ii) DIC incubation. The three remaining jars amended with ^13^C^15^N-BCAA were used as seawater control incubations. All incubation jars (n = 9) were randomly distributed in a chest-type orbital shaker incubator (Thermoline Scientific), which was set to 80 rpm and 28.5°C, matching the *in situ* seawater temperature. All jars were kept in the dark throughout the experiment. Oxygen concentrations were monitored throughout the experiments via oxygen sensor spots and the FireStingGo2 pocket oxygen meter (PyroScience GmbH). Pure oxygen gas was added with a syringe and needle to the headspace of the incubation jars containing sponge explants to maintain 100% oxygen saturation throughout the 24h incubation period.

Samples of all jars (n = 9) were collected at five time points throughout the incubation experiments via the sampling port septum of the incubation jars using a needle and a syringe, and sterile filtered (PES filter, 0.2µm). The collected, filtered incubation water samples were used to determine the nitrification activity (stored at - 20°C), the BCAA concentrations (BCAA experiment only, stored at −80°C), the ^13^C-DIC label percentage (stored at 4°C), and NH_4_^+^ concentrations (stored at −20°C, see EDF 4). After the last sampling time point, the wet weight (ww) of each sponge explant was measured. Sponge explants of the BCAA experiment were washed three times for 20 min in 0.2 µm filter-sterilized seawater (Sterivex, Merck Pty Ltd) at 80 rpm at 28.5°C to remove excess isotopically labeled BCAA from the tissue. The tissue was then briefly rinsed with 0.2 µm filter-sterilized seawater and subsampled for DNA extractions, CARD-FISH-NanoSIMS imaging, bulk enrichment with stable isotopes, and sponge pore water concentrations of BCAA. The remaining incubation water in the incubation jars (approximately 760 mL) was individually filtered onto 0.2 µm Sterivex filters for DNA extractions (n = 3 for BCAA seawater control samples, n = 1 for sponge incubations).

### ^13^C-DIC measurements on GC-IRMS

^13^C-DIC production from ^13^C^15^N-labeled BCAA was assessed by conversion of all DIC to CO_2_ by acidification and subsequent measurement of ^13^C/^12^C ratios in CO_2_ using gas chromatography - isotope ratio mass spectrometry (GC-IRMS). Briefly, 12 ml glass vials (Labco Exetainer) were amended with 25 µl 20% phosphoric acid, closed airtight with a screw cap containing a septum, and flushed with He for 3 minutes. Then, 0.5 ml aliquots of seawater samples were injected into the closed, flushed glass vials, carefully swirled to mix with phosphoric acid, and incubated overnight before measurement of the CO_2_ released into the headspace using a gas preparation system connected to the GC-IRMS (GasBench II connected to Delta V Advantage IRMS, Thermo Scientific, Vienna, Austria). ^13^C/^12^C ratios in CO_2_ were calibrated against reference CO_2_ standards and freshly prepared standards of NaHCO_3_ (natural abundance).

### BCAA concentrations in sponge “pore-water” and incubation water samples

Valine, leucine, and isoleucine concentrations were measured in sponge pore water samples (n = 9) and in incubation water samples (n = 30) of the BCAA incubation experiment. Sponge pore water was recovered by placing fresh sponge fragments into 0.5 ml plastic centrifuge tubes with perforated bottoms (5 holes punched with a 23 G needle). The perforated tubes were placed into 2 ml plastic collection tubes, and *I. basta* pore water was collected via centrifugation at 1,000 g for 1 min. The freshly collected incubation water was filtered through a 0.2 µm syringe filter (CHROMAFILL). All collected samples were immediately frozen and stored at −80 °C until further analysis.

Prior to analysis, pore water samples (n = 9) were diluted 1:10 in artificial seawater (per L: 26.4 g NaCl, 6.8 g MgSO_4_*7H_2_O, 5.7 g MgCl_2_*6H_2_O, 1.47 g CaCl_2_*2H_2_O, 0.66 g KCl, 0.2 g KH_2_PO_4_, 0.19 g NaHCO_3_) and the remaining debris was pelleted by centrifugation at 10,000 g for 5 min. BCAA concentrations were quantified against the respective standards (0.0625 to 2 µM) prepared in artificial seawater. Samples and standards were derivatized with the AccQ-Tag Ultra Derivatization Kit (WatersTM) and subsequently measured on a UPLC-Orbitrap mass spectroscopy system ^70^ with modifications to the UPLC settings. In brief, samples were measured using a UPLC Ultimate 3000 system (Thermo Fisher Scientific, Bremen, Germany) linked to an Orbitrap Q Exactive HRMS system (Thermo Fisher Scientific) with an electrospray ionization (H-ESI) source. Chromatographic separation was conducted on a Water AccQ-Tag Ultra C18 column (2.1 × 100 mm, 1.7 µm particles; Milford, MA, USA) with an ACQUITY column inline guard filter (2.1 mm, 0.2 µm; Milford, MA, USA) at 55 °C column temperature. The mobile phase consisted of 0.001% formic acid in MilliQ (eluent A) and 0.001% formic acid in acetonitrile (eluent B). The following solvent gradient was applied: 0–0.5 min kept constant at 0.1% B, 0.5–2.5 min increased to 5% B, 2.5–8 min increased to 20% B, 8–8.25 min increased to 90% B, 8.25–11 min maintained at 90% B, 11–11.2 min decreased to 0.1% B, and 11.2–4.5 min maintained at 0.1% B for column re-equilibration. The flow rate was set to 0.4 ml min^−1^ and the injection volume was 1 µL. The Orbitrap Q Exactive HRMS system was operated using full-mass scan mode (m/z 150–1000) in the ESI positive mode. The automatic gain control (AGC) target values were set to 3 × 106 and the resolution was set to 70,000. The lock mass correction was set to 214.0896 (contaminant from plasticizer). The spray voltage was 3.5 kV, while the capillary temperature was set to 300 °C. The sheath gas consisted of 35 arbitrary units, and the auxiliary gas of 15 arbitrary units.

### Bulk nitrification rates and ^15^N nitrite formation from BCAA

Bulk nitrite and nitrate production in the incubations was analyzed photometrically using the Griess protocol ^71,72^. Measured concentrations were converted to nitrite (or nitrate) production rates per sponge wet weight by accounting for the removal of incubation water aliquots during sampling.

^15^N-nitrite production from ^13^C^15^N-BCAA was analyzed by conversion of nitrite to nitrous oxide using azide ^73^ and subsequent measurement of ^15^N-N_2_O to ^14^N-N_2_O ratios using GC-IRMS. Briefly, 500 µL sample were incubated at 37°C and 100 rpm shaking with 100 µl of 1 M NaN_3_ in 5% acetic acid and 500 µL 1M HCl for 24h. The reaction was stopped by injecting 100 of µL 6 M NaOH. For ^15^N-N_2_O measurement, the samples were purged with helium gas at 10 ml min^-1^, passed through a magnesium perchlorate and ascarite trap to remove water vapour and CO_2_, and cryo-focused in two consecutive liquid nitrogen traps (1^st^ trap stainless steel and 2^nd^ trap fused silica, Finnigan Precon, Thermo Fisher). The sample gas was released at room temperature and transferred through a Nafion trap into a GC-column (GS-G, Gasbench II head space analyzer, Thermo Fisher) to separate N_2_O from other gases and finally injected into the ion source of the IRMS instrument (Finnigan Delta Advantage V, Thermo Fisher).

### 16S rRNA gene amplicon sequencing and analysis

DNA from (i) field-control sponge explants (n = 4), (ii) sponge explants from the BCAA incubation experiment (n = 3) and the DIC incubation experiment (n = 3), (iii) and the 0.2 µm Sterivex filter (Merck Pty Ltd) from the seawater control jars (n = 3) and sponge incubation jars (n = 2) from the BCAA incubation experiment was extracted using the DNeasy PowerSoil Pro Kit (Qiagen). Blank DNA extractions were included to account for possible contamination. The V4 region of the 16S rRNA gene was amplified and sequenced on an Illumina MiSeq (2 x 300bp) at the Joint Microbiome Facility (JMF) of the Medical University of Vienna and the University of Vienna, using the primer pair 515F 5′-GTG YCA GCM GCC GCG GTA A-3′ ^74^ and 806R 5′-GGA CTA CNV GGG TWT CTA AT-3ʹ ^75^, as previously described ^76^. Demultiplexing was performed with the python package demultiplex (Laros JFJ, github.com/jvlaros/demultiplex), allowing one mismatch for barcodes and two mismatches for linkers and primers. Amplicon sequence variants (ASVs) were inferred using the DADA2 R package v1.32 ^77^, applying the recommended workflow (https://f1000research.com/articles/5-1492). FASTQ reads 1 and 2 were trimmed at 220 nt and 150 nt, with allowed expected errors of 2. ASVs were classified using DADA2 against the SILVA database SSU Ref NR 99 release 138.1. ASVs classified as chloroplasts and mitochondria were removed from the ASV table. Singletons (ASVs that occur only once) were removed from the ASV table for the subsequent data analysis. The relative abundances of the ASVs originating from the three dominant *I. basta* symbionts, together with the other bacterial and archaeal ASVs were calculated and visualized in R ^78^ using the packages phyloseq ^79^, dplyr ^80^, and ggplot2 ^81^.

### ^13^C and ^15^N enrichment in bulk sponge tissue

The ^13^C and ^15^N enrichment in the bulk tissue of sponge explants from the DIC and BCAA incubation experiments and the natural abundance of these isotopes in field-control explants were measured by elemental analyzer-IRMS (EA -Isolink coupled via ConFlo IV interface to Delta V Advantage IRMS, Thermo Scientific). Prior to analysis, the subsampled fragments of sponge explants (n = 9) were freeze-dried and ground to a fine powder in a Retsch MM200 ball mill (Retsch, Hanau, Germany) for 5 min. Samples for ^13^C enrichment measurements were decalcified with 0.4 M HCl overnight to remove inorganic carbon and dried at 65 °C for 48 hours. All samples were subsequently weighed (between 0.5 - 1 mg) into tin capsules and stored dry until measured. The EA-IRMS system was calibrated for carbon and nitrogen content and isotope composition using lab standards (sugar-amino acid mixes) referenced against international standards (USGS40, USGS41a, USGS64). Heavy stable isotope abundances of carbon and nitrogen were measured in untreated (natural abundance) samples and in isotopically enriched ones as atom% ^13^C and atom% ^15^N, and the atom% of treated samples were then contrasted against control samples by calculating the atom percent enrichment (APE) of ^13^C and ^15^N.

### CARD-FISH - NanoSIMS of sponge tissue sections

Subsamples of sponge explants of both labeling experiments (BCAA and DIC) and a natural abundance control sample of an untreated *I. basta* individual were fixed in 4% paraformaldehyde (ProSciTech) overnight at 4°C, rinsed three times with ice-cold 1x phosphate-buffered saline (PBS), and subsequently stored in PBS:ethanol (1:1) at −20°C. Thin sections of sponge tissue were prepared at the Histology Facility of the Vienna Biocenter (Vienna, Austria). In brief, fixed samples were washed for 1 h in 1x PBS and stored overnight at 4°C in 30% sucrose to preserve tissue morphology. Samples were subsequently transferred to 50% sucrose:Tissue-Tek O.C.T (Optimal Cutting Temperature) compound embedding medium (Sakura) for embedding, and stored overnight at 4°C. The embedded samples were transferred to isopentane and frozen in liquid nitrogen. Embedded tissue blocks were kept either on dry ice or at −80°C until 5 µm-cryosections were prepared and transferred onto Indium Tin Oxide (ITO) wafers.

Cryomedium residues were removed by submerging the wafers for 3 min each in 50%, 80%, and 96% ethanol. Subsequently, samples were dried at room temperature. To enable correlative imaging of fluorescent visualization of the sponge symbionts via catalyzed reporter deposition fluorescence *in situ* hybridization (CARD-FISH) and their single-cell isotope incorporation via nanoscale secondary ion mass spectrometry (NanoSIMS), regions of interest were marked using a laser microdissection microscope (Leica LMD 7000, Germany).

CARD-FISH was performed following Pernthaler, et al. ^82^, using probes Arch915 ^83^, AlfD729 ^84^, and GamD1137 ^13^, together with newly designed helper probes (Supplementary Table 1), with some modifications. For all CARD-FISH experiments, negative controls using probe NonEUB ^85^ were included. Prior to CARD-FISH, sponge tissue sections were encircled using a pap-pen (Kisker, MKP-1) to avoid loss of buffers during the CARD-FISH protocol. The samples were permeabilized using lysozyme (10 mg ml^-1^ in 0.05M EDTA, 0.1M Tris*HCl, pH 8, 60 min at 37°C). After washing in deionized water twice, the samples were additionally permeabilized using HCl (0.2M, 10 min, room temperature), again washed twice in deionized water, and dried. Endogenous peroxidases were inactivated using a freshly prepared 0.3% H_2_O_2_ solution in 1x PBS (30 min, room temperature), samples were again washed in deionized water, and dried. We performed two consecutive rounds of hybridization (each 3 h, 46°C, at the respective optimal formamide concentration, see Supplementary Table 1) and signal amplification (each 30 min, 46°C), using Alexa488-labeled tyramides with probe Arch915 and Alexa594-labeled tyramides with probe AlfD729. Between each round of hybridization, the previously used probes were inactivated by incubation in 0.01M HCl (10 min, at room temperature), followed by washing in deionized water twice.

Following CARD-FISH, samples were counterstained using DAPI (1 µg ml^-1^, 10 min, room temperature), washed twice in deionized water, and let dry, before embedding in CitiFluor AF1 (EMS). CARD-FISH signals of AOA symbiont *Ca*. Nitrosospongia ianthellae and alphaproteobacterial symbiont Ca. *Luteria ianthellae*, as well as LMD markings were visualized using a confocal laser scanning microscope (SP7, Leica, Germany, equipped with a white light laser). The gammaproteobacterial symbiont *Ca*. Taurinisymbion ianthellae cells were identified based on DAPI signals showing their characteristic rod-shaped morphology, which we confirmed in separate CARD-FISH experiments using the specific probe GamD1137 ^13^. To capture symbionts within the entire volume of the sponge sections, z-stacks were recorded and projected into one plane (maximum projection algorithm). After visualization, the embedding medium was removed from the sections by washing in deionized water three times for 5 min each.

NanoSIMS measurements were carried out using a NanoSIMS 50L instrument (Cameca, Gennevilliers, France) at the Large-Instrument Facility for Environmental and Isotope Mass Spectrometry, University of Vienna. Prior to data acquisition, analysis areas were pre-conditioned by rastering with a high-intensity, defocused Cs⁺ ion beam using extreme low ion impact energies (EXLIE) deposition procedure reported previously ^86^, shortly: high energy (HE, 16 keV) at 100 pA beam current to a fluence of 5 × 10^14^ ions per cm^2^; EXLIE (50 eV) at 400 pA beam current to a fluence of 5 × 10^16^ ions per cm^2^; HE to an additional fluence of 2.5 × 10^14^ ions per cm^2^. The multicollection detector assembly was configured to enable parallel detection of the following secondary ions: ^12^C^12^C^-^,^12^C^13^C^-^,^12^C^14^N^-^,^12^C^15^N^-^, ^31^P^-^, and ^32^S^-^. The mass spectrometer was tuned to achieve a mass resolving power (MRP) >9,000 (as defined by Cameca) to ensure accurate separation of C_2_^-^ and CN^-^ secondary ions. Data were acquired as multilayer image stacks by sequential rastering of a finely focused Cs⁺ primary ion beam (∼1 pA), achieving a lateral resolution of ∼80 nm. Scanning areas ranged from 25 × 25 µm^2^ to 50 × 50 µm^2^ at 512 × 512 pixel resolution, with a per-pixel dwell time of 1.5 ms per pixel per cycle. Between 38 to 47 cycles were acquired per scanning area.

NanoSIMS images were processed using Look@NanoSIMS (version 24-12-20) implemented in MATLAB R2024b (academic license). Image stacks (cycles) were aligned to correct for positional drift due to room temperature fluctuations, primary beam instability, or stage movement. Secondary ion intensities were corrected on a per-pixel basis for detector dead time and quasi-simultaneous arrival effects using sensitivity correction factors of 1.06 for C_2_^-^, 1.05 for CN^-^, and 1.10 for all other ions. All measurement cycles were accumulated, and regions of interest (ROIs), corresponding to individual microbial cells or sponge nuclei, were manually defined based on phosphorus ion distribution maps in correlation with CARD-FISH and DAPI signals. Note that because ROIs were defined using all accumulated cycles per measurement area, it is possible that some ROI enrichments are under- or overestimated when secondary ions from tissue below or above microbial cells were ablated. We did not take any dilution effects from the CARD-FISH procedure into account for our calculations, as they differ depending on the CARD-FISH protocol used, cell types, and growth stages ^87–89^, which cannot be constrained in environmental samples. As such, the reported growth rates, doubling times, and amounts of freshly fixed carbon and nitrogen are considered conservative.

Carbon isotope composition (expressed as ^13^C-atom%) was calculated from C_2_^-^ secondary ion images using Eq. 1:

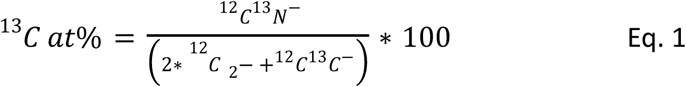

Similarly, nitrogen isotope composition (^15^N-atom%) was derived from CN^-^ secondary ion images using Eq. 2:

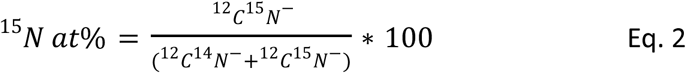

Only ROIs with an associated Poisson error <5% (for carbon or nitrogen isotopic composition, respectively) were considered for further calculations and used for visualization.

Single-cell carbon-based growth rates (GR) were calculated following Martinez-Perez, et al.^90^ using Eq. 3

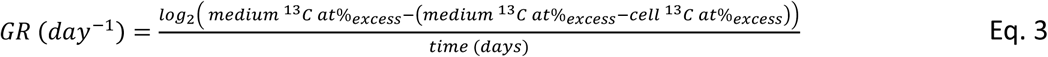

Where time is the incubation time in days. Medium ^13^C at% _excess_ was obtained for DIC incubations using equation 4:

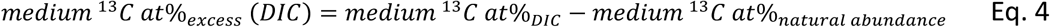

Medium 13C at% _excess_ for BCAA incubations was determined using equation 5:

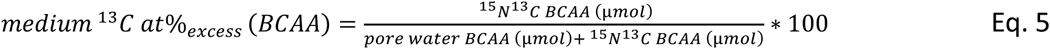

Cell ^13^C at%_excess_ was obtained by subtracting the mean ^13^C at% of all single symbiont cells in natural abundance sponge sections (1.06 ^13^C at%) from single cell ^13^C at% values determined in ^13^C-DIC and ^15^N^13^C-BCAA incubated samples.

Doubling times were inferred from Eq. 6:

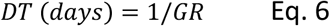

The amount of newly assimilated carbon (and analogously for nitrogen) from BCAA was estimated following Krupke, et al. ^91^ using Eq. 7:

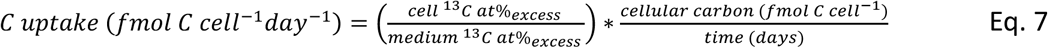

Cellular carbon contents were estimated following Khachikyan, et al. ^92^, using ROI dimensions from nanoSIMS data and Eq. 8 and Eq. 9:

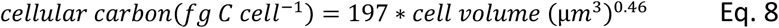

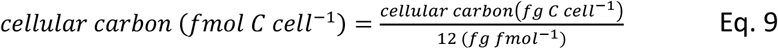

Where cell volume (in µm^3^) was estimated from nanoSIMS ROI dimensions, assuming prolate sphere shapes using Eq. 10 ^93^:

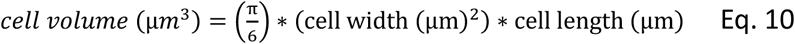

Cellular nitrogen content was estimated using the Redfield ratio of C:N = 6.625 (Eq. 11):

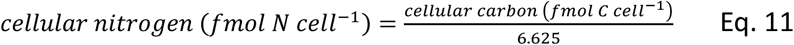

Note that we assumed that the sponge symbionts are exposed to the concentrations of experimentally added isotope tracers inside the sponge holobiont, i.e., that the DIC concentrations and ^13^C-isotope enrichment were identical in the incubation seawater (medium) and inside the sponge. For BCAA, we assumed that the experimentally added ^13^C^15^N-BCAA was available to the sponge symbionts at equal concentrations as supplied to the incubation seawater. Furthermore, we assumed that the sponge symbionts are additionally exposed to BCAA concentrations of natural abundance measured in sponge pore water. Whether these assumptions hold true requires additional research; however, we assume that if the sponge holobiont indeed reduces the availability of stable isotope-labeled compounds for its symbionts by providing an uptake barrier, this effect would be similar for both BCAA and DIC.

## Acknowledgements

For the purpose of open access, B.G. has applied a CC BY public copyright license to any author-accepted manuscript version arising from this submission. We thank Stefanie Imminger and Arno Schintlmeister for their help and advice on the NanoSIMS sample preparation. We also thank Martina Mayer for her support in measuring ^15^N-nitrite enrichment, Emma Marangon for her help during the sponge incubations, Raimund Schubert for the construction of our sponge-incubation jars, and Holger Daims for the critical feedback and discussions. We further thank the SeaSim staff, the RV Cape Ferguson crew, and the RV Apollo crew for their help with the sponge collection and aquaria maintenance, the AIMS Analytical Technology Team for measuring ammonium concentrations, the Vienna BioCentre Histology Facility for preparing cryosections of the sponge tissue and the CeMESS Technical Assistant Team, Microscopy Team, JMF Laboratory Team and the Life Science Compute Cluster for their support throughout the project. We further acknowledge the traditional owners of land and sea country, the Bindal, Manbarra, and Wulgurukaba people, where parts of the research have been conducted, and we respect their elders past, present, and emerging.

