## Extended data figures and supplementary Table for "Branched-chain amino acid assimilation promotes mixotrophy of ammonia-oxidizing archaeal sponge symbionts"

**Content:**

- 8 Extended Data Figures (EDF)
- 1 Supplementary Table

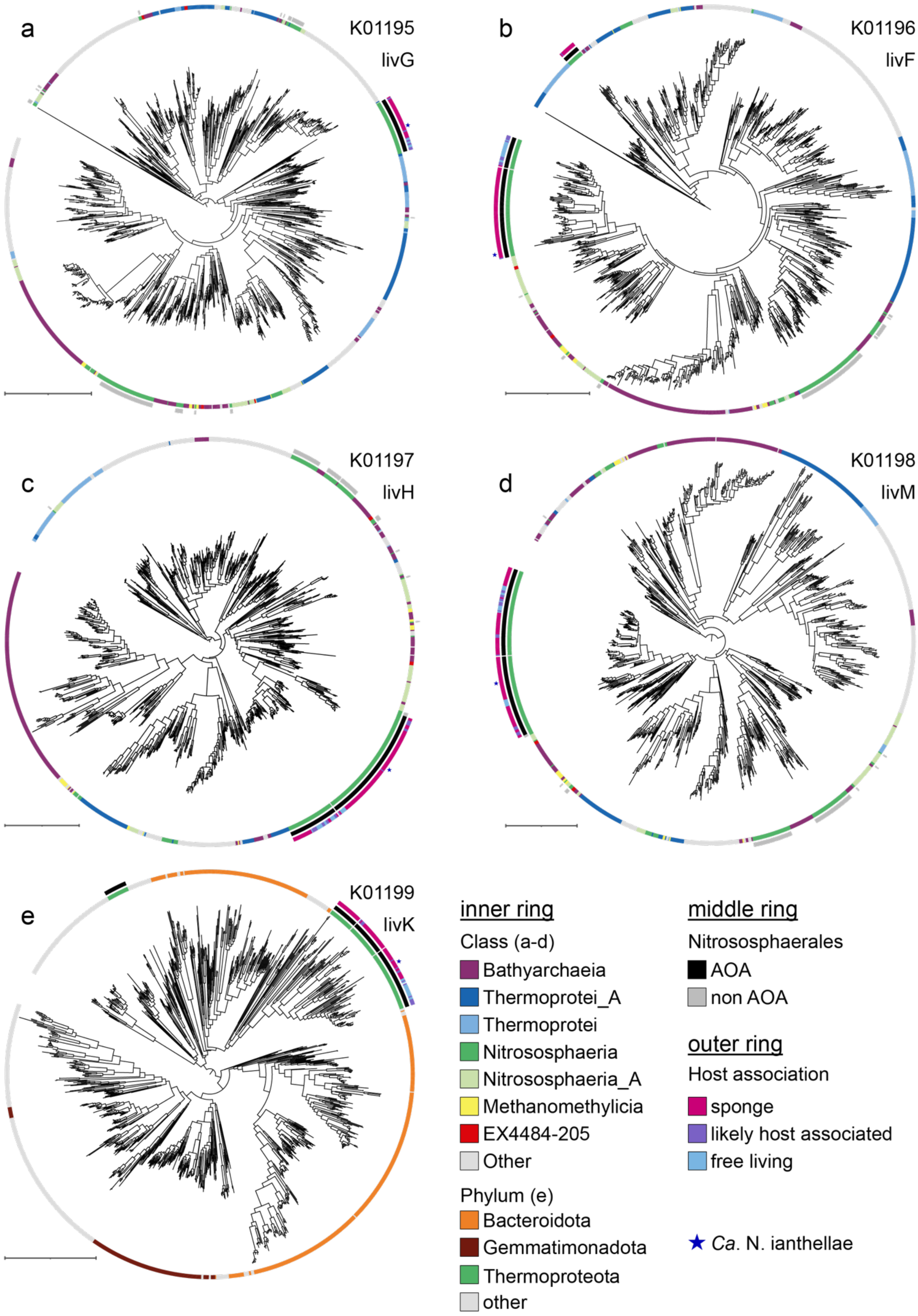


**EDF 1. Amino acid phylogenies of the phylogenetic context of the BCAA transporter subunits in ammonia-oxidizing archaea (AOA).** The taxa included in the phylogenies were chosen based on a

phylogenetic analysis of the full datasets of each subunit (see methods). The phylogenetic neighborhood of the ATP-binding subunits *livG* (a) and *livF* (b), and the transmembrane subunits *livH* (c) and *livM* (d), largely consists of conserved taxa within the archaeal *Thermoproteota* phylum that also includes the *Nitrososphaerales*. While this suggests a shared ancestry, the topology of the four phylogenies is not consistent, making it impossible to determine the specific donor lineage based on these analyses. In contrast, the phylogenetic neighborhood of the AOA substrate binding subunit *livK* (e) consists primarily of sequences of the bacterial phylum *Bacteroidota*, indicating a different ancestry than the rest of the complex. Inner colored rings around the phylogenies depict taxonomic affiliation at the class (a-d) or phylum (e) level. The middle colored ring indicates sequences from genomes in the *Nitrososphaerales* order, with AOA in black and other *Nitrososphaerales* in grey. The outer colored ring indicates whether the source genome of a sequence is host-associated based on the source data of the genome and the metagenomes in which it is detected. The position of the *Candidatus* Nitrosospongia ianthellae sequences is indicated with a blue star. The scale bar at the bottom right of each phylogeny indicates one substitution per site.

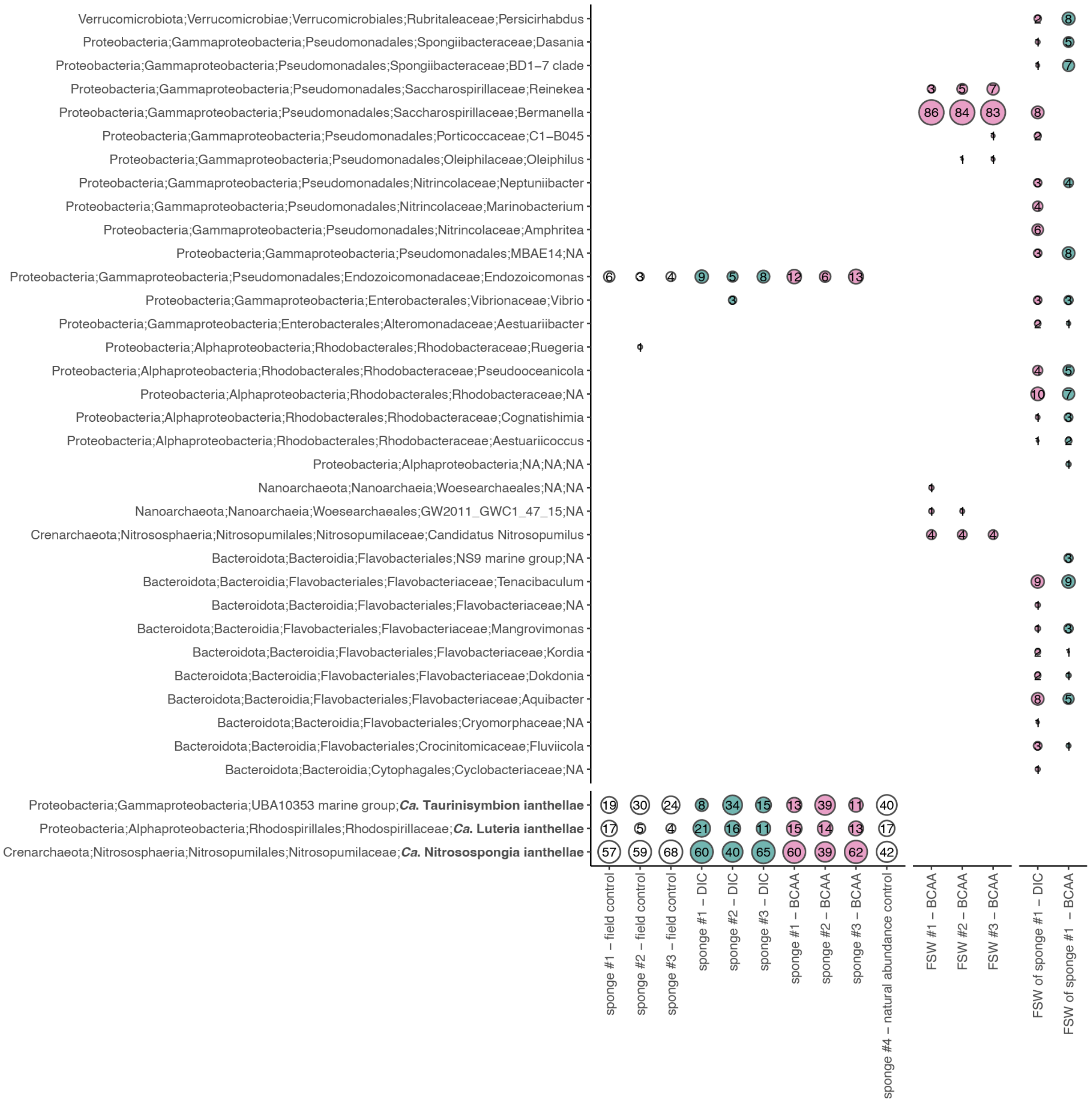


**EDF 2. Microbial community composition of *Ianthella basta* and seawater samples.** The relative abundance of bacterial and archaeal microbiome members in *I. basta* individuals was determined using 16S rRNA gene amplicon sequencing. The microbiome composition of the sponge explants remained stable throughout the stable isotope incubation experiments (DIC = $^{13}$C-labeled dissolved inorganic carbon incubation experiment, BCAA = $^{13}$C$^{15}$N -labeled branched-chain amino acids incubation experiment). Sponge-explants were subsampled prior to the experiments (field control) and at the end of the 24h incubation experiments. The microbial community associated with the filtered seawater control samples, collected at the end of the 24h incubation period, of the $^{13}$C$^{15}$N-labeled BCAA incubation experiment (FSW BCAA, pore size for filtration 0.22 mm; EDF 4) and the incubation water of the sponge incubations with labeled DIC and BCAA (FSW of sponge #1; EDF 4 and 5) varied substantially from the *I. basta* microbiome. In the filtered seawater BCAA control jars, we

observed growth of putative heterotrophs belonging to the genus *Bermanella* (Pinhassi et al. 2009), which either passed through the filter pores (0.22 µm) or were present in the incubation jars/room during experimental setup, which was done under non-sterile conditions. The growth of these putative heterotrophs can also explain the consumption of BCAA in the filtered seawater incubation without sponge explants (EDF 4). The relative abundances of the three dominant *I. basta* symbionts are displayed at the amplicon sequence variant (ASV) level; the relative abundances of the other microbiome ASVs are summarized at the family level.

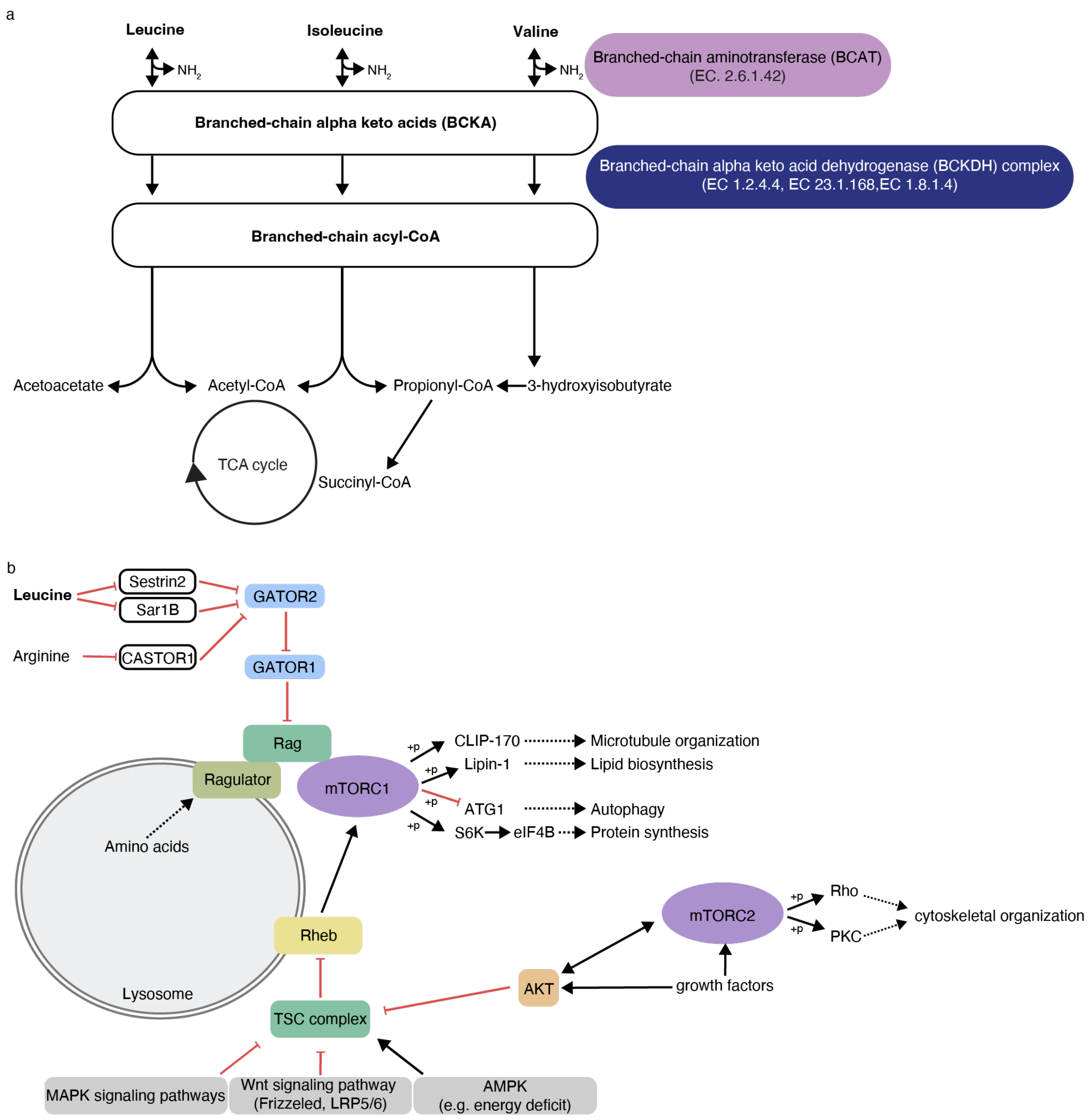


**EDF 3. Genomic capacity of *Ianthella basta* to utilize branched-chain amino acids (BCAA).** a) The *I. basta* (GCA_964019415.1) sponge genome encodes for the complete branched-chain amino acid (BCAA) degradation pathway. The first step is the reversible transamination of BCAA to its respective branched-chain alpha keto acids (BCKA). The second step is the oxidative decarboxylation of the BCKA to branched-chain acyl-CoA derivatives, which is catalyzed by the branched-chain alpha-keto acid dehydrogenase (BCKDH) complex. Succinyl-CoA and acetyl-CoA are the end products of the valine, leucine, and isoleucine catabolism. Succinyl-CoA and acetyl-CoA serve as substrates for the tricarboxylic acid (TCA) cycle. b) BCAA, especially leucine, are key regulators of the mechanistic target of rapamycin complex 1 (mTORC1), a central protein complex of eukaryotic cells. Leucine can modulate the mTORC1 complex and thus stimulate microtubule organization, lipid biosynthesis, and protein synthesis, but inhibits autophagy. The *I. basta* genome encodes all genes of the mTORC1 complex but lacks homologous genes encoding for the leucin sensing protein Sestrin2 and Sar1B, and

the arginine sensing protein CASTOR1. The lack of homologous leucin-sensing genes in *I. basta* can be due to the non-closed status of the genome (79% complete), too low homology to the previously described leucin-sensing genes, or yet unknown genes and/or regulators of mTORC1. Future studies will need to address the role of BCAA on mTORC1 in sponges and search for potential candidate genes based on homologous structures. KEGG Orthology (KO) assignment and KEGG pathway reconstruction were performed using GhostKOALA (Kanehisa et al. 2016). The pathway maps were generated with KEGG mapper and modified in Adobe Illustrator.

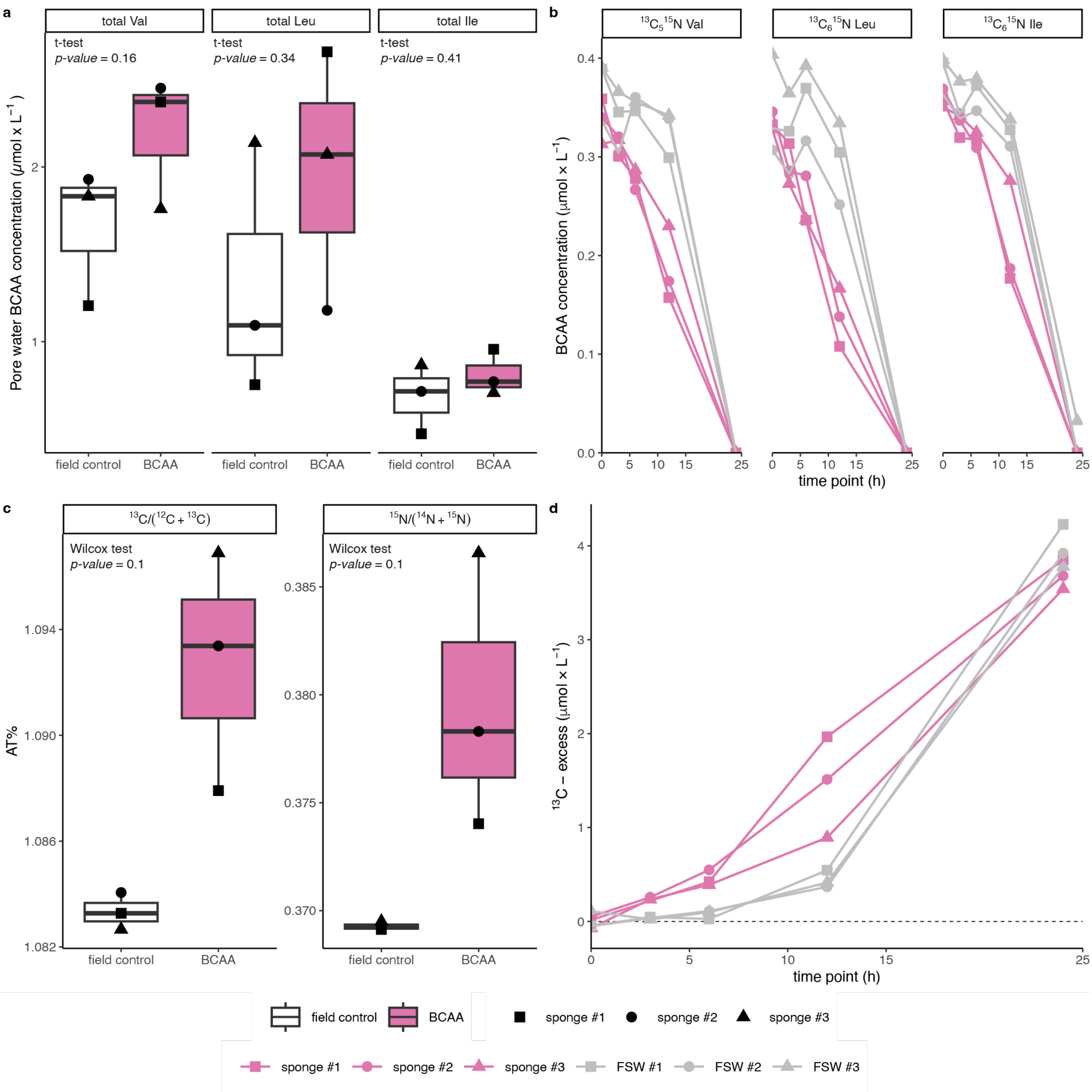


**EDF 4. Bulk BCAA concentration in and turnover by the *Ianthella basta* holobiont** a) Total branched-chain amino acid (BCAA) concentrations (i.e., valine, leucine, and isoleucine) in readily extractable pore water from *I. basta* holobiont samples (field control = sponge pore water samples collected prior to incubation, BCAA = sponge pore water samples collected after 24h BCAA incubation experiment). Boxplots depict the 25–75% quantile range, with the centre line depicting the median (50% quantile); whiskers encompass data points within 1.5× the interquartile range. b) Depletion of isotopically labeled BCAA in the incubation water over the 24h incubation period. c) $^{13}C$ and $^{15}N$-enrichment in *I. basta* holobiont biomass samples (field control = sponge biomass samples collected prior to incubation, BCAA = sponge biomass samples collected after 24h BCAA incubation experiment). Boxplots depict the 25–75% quantile range, with the centre line depicting the median (50% quantile); whiskers encompass data points within 1.5× the interquartile range. d) The respiration of isotopically

labeled BCAA results in an excess of $^{13}C$-labeled $CO_2$ in the incubation water over the course of the 24h experiment. Note that also in incubations containing only filtered seawater (no sponge explants), added BCAA are depleted over the course of the incubation, and largely respired to $CO_2$. This can be explained by the presence and growth of some heterotrophic cells that are capable of respiring BCAA, which either passed through the filter pores (0.22 µm) or were present in the incubation jars/room during experimental setup, which was done under non-sterile conditions. Putative heterotrophs growing during incubation belong to the genus *Bermanella* (Pinhassi et al. 2009), which we detected differentially enriched in filtered seawater incubations without sponge explants (EDF 2).

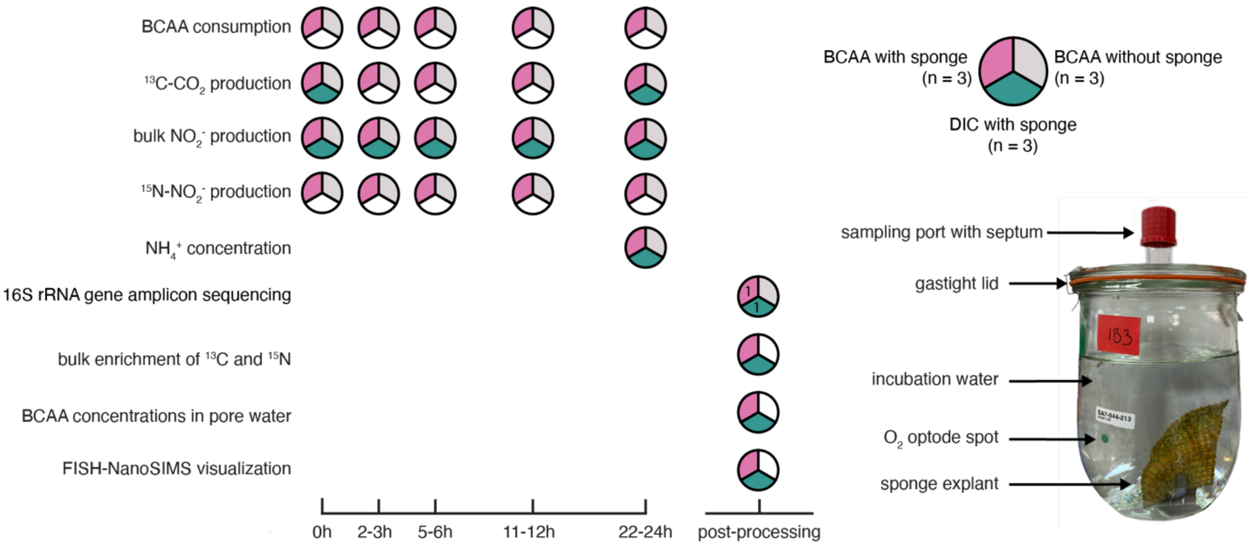


**EDF 5. Sampling scheme and experimental setup for the stable isotope incubation experiments.** The incubation jars were filled with filtered seawater and amended with a $^{15}N^{13}C$-labeled branched-chain amino acid (BCAA) mix containing equimolar concentrations of valine, leucine, and isoleucine, or with $^{13}C$-dissolved inorganic carbon (DIC). Incubation water samples for BCAA consumption, $^{13}C$-$CO_2$ concentrations, bulk $NO_2^-$ , $^{15}N$-$NO_2^-$ , and $NH_4^+$ concentrations were collected at regular intervals via the sampling port throughout the incubation period (up to 24 hours). At the end of the incubation experiments, the sponge explants were subsampled for (i) 16S rRNA gene amplicon sequencing, (ii) measurement of bulk enrichment of $^{13}C$ and $^{15}N$ in the sponge tissue, (iii)extraction of pore water samples, and (iv) fixation for FISH-NanoSIMS. Incubation jars remained closed throughout the incubation period to avoid exchange of the headspace with the atmosphere.

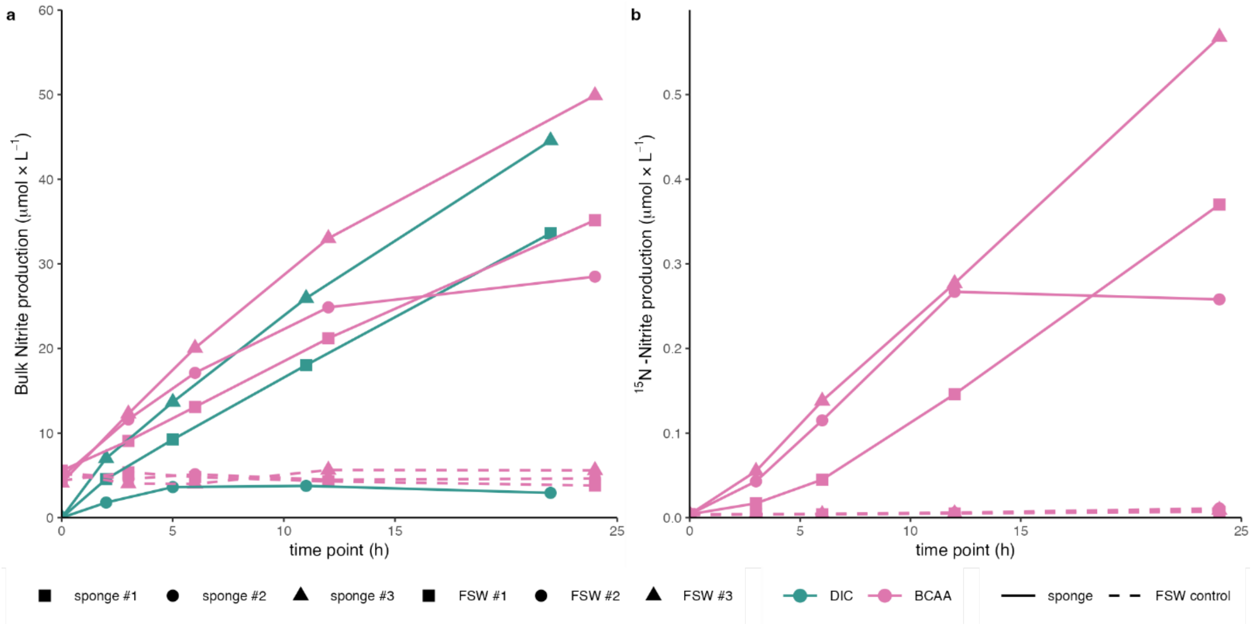


**EDF 6. Nitrification activity of the *Ianthella basta* sponge holobiont and seawater control jars.** a) Bulk nitrite ($NO_2^-$) production (in µmol per L) in sponge and seawater control jars during DIC and BCAA incubation experiments. b) $^{15}$N-nitrite ($NO_2^-$) production (in µmol per L) in sponge and seawater control jars during the BCAA incubation experiment. Bulk and $^{15}$N-Nitrite production normalized to the sponge wet weight are shown in Figure 3.

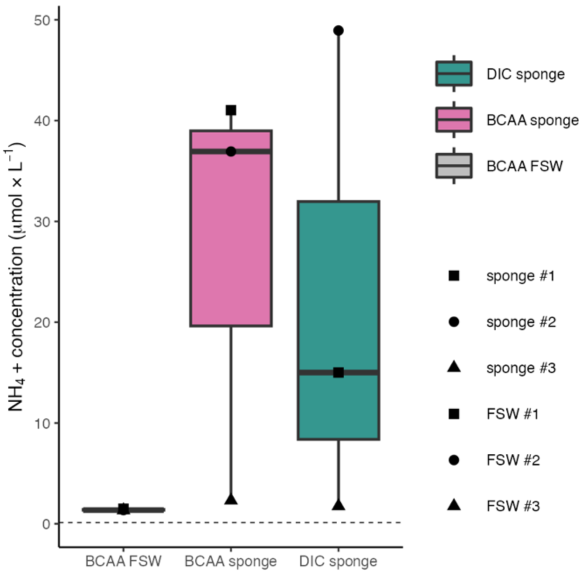


**EDF 7.** Ammonium ($NH_4^+$) concentration in the seawater control jars filled with sterile filtered seawater (FSW) and in the sponge jars in the branched-chain amino acid (BCAA) incubation and dissolved inorganic carbon (DIC) experiments was measured at the end of the incubation experiments. Boxplots depict the 25–75% quantile range, with the centre line depicting the median (50% quantile); whiskers encompass data points within 1.5× the interquartile range. The dashed line represents the *in situ* seawater $NH_4^+$ concentration of the field sampling site.

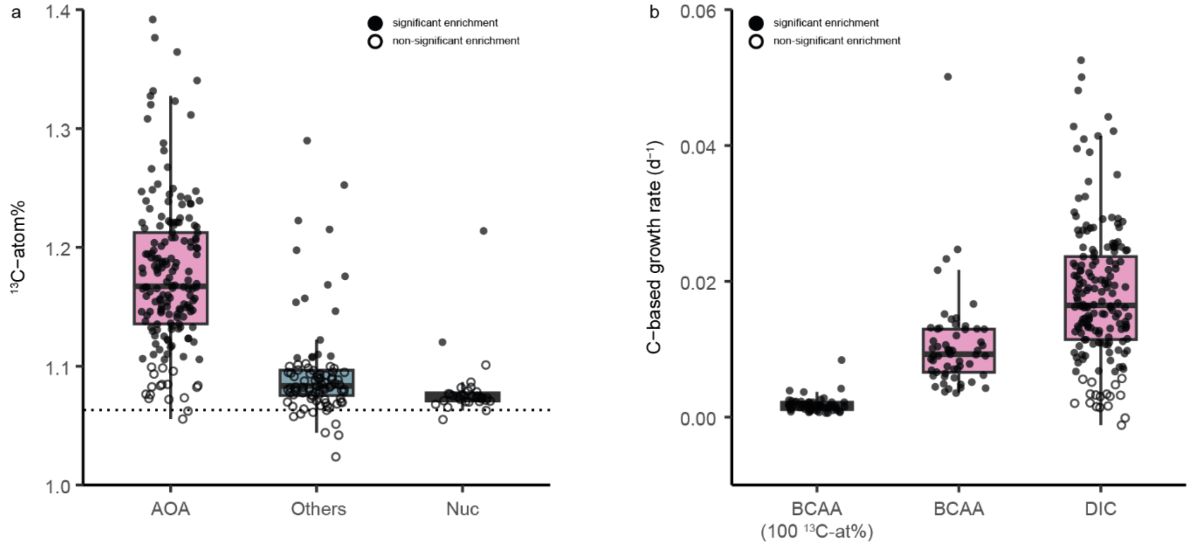


**EDF 8.** Single cell $^{13}C$-enrichment from $^{13}C$-DIC and comparison of C-based AOA growth rates in incubations with $^{13}C^{15}N$-BCAA and $^{13}C$-DIC. a) Single cell $^{13}C$-enrichment of AOA (*Ca.* N. ianthellae) and other microbial symbionts of *I. basta* (Others) and sponge nuclei (Nuc). Dotted line indicates $^{13}C$ natural abundance at% as measured for symbiont cells in non-incubated sponge biomass (1.06 $^{13}C$ at%). AOA are significantly more enriched compared to the other microbial symbionts (one-sided Mann–Whitney U test, W = 14,007, $p < 2.2 \times 10^{-16}$) b) Carbon-based growth rates of single AOA cells from $^{13}C^{15}N$-BCAA, assuming that symbionts were either exposed to only fully labeled $^{13}C^{15}N$-BCAA (BCAA (100 $^{13}C$-at%)), or that symbionts were exposed to experimentally added $^{13}C^{15}N$-BCAA and BCAA of natural abundance present in the sponge pore water (BCAA). Growth rates inferred from $^{13}C$-DIC assimilation are also depicted. Boxplots depict the 25–75% quantile range, with the centre line depicting the median (50% quantile); whiskers encompass data points within 1.5× the interquartile range.

**Supplementary Table 1.** Oligonucleotide probes used in this study.

| Oligo ID | Formamide concentration | Sequence | Helper Sequence (designed in this study) |
|---|---|---|---|
| Arch915 (Stahl and Amann 1991) | 30% | GTG CTC CCC CGC CAA TTC CT | - |
| AlfD729 (Moeller et al. 2019) | 25% | CGG ACC TGG CGG CCG CTT | GCT ACT GGT GTT CTT CCC AAT CTC TAC GAA TTC CAC; CAC GCT TTC GCG CCT CAG CGT CAG T; CTG GGA GTT CCG CCG TCA TTT TCC G |
| GamD1137 (Moeller et al. 2019) | 25% | CTC AAA GTC CCC GCC ATT | GCG CTG GCA ACT AAG GAC AAG; CCG GTT TGT CAC CGG CAG TC |
| NonEUB (Wallner et al. 1993) | 30% | ACT CCT ACG GGA GGC AGC | - |
